# The glutamine transporter Slc38a1 is widely expressed in the embryonic neurogenic niches and impacts neuronal volume, survival, and morphology

**DOI:** 10.64898/2025.12.16.694571

**Authors:** Daxin Wang, Marivi Nabong Moen, Diana Domanska, Muhammad Zahoor, Wenjing Cai, Ragnhild Elisabeth Heimtun Paulsen, Tor Paaske Utheim, Frode Lars Jahnsen, Jon Storm-Mathisen, Farrukh Abbas Chaudhry

**Author notes:** **Corresponding author:** Farrukh Abbas Chaudhry, Department of Molecular Medicine, Institute of Basic Medical Sciences, University of Oslo, Norway.

## Abstract

The amino acid glutamine, and its derivatives glutamate and GABA, are pivotal for neurogenesis. However, how glutamine is mechanistically supplied to the cells in the neurogenic niches and its broader impact on neurodevelopment, remain poorly characterized. The Solute carrier family 38 member 1 (Slc38a1) exhibits high affinity for glutamine and accumulates glutamine in select cells. We investigated whether Slc38a1 is essential for neurogenesis and whether it has an impact on brain structure and function. Slc38a1 mRNA transcript and protein are widely expressed in the embryonic brain, including in the dorsal pallium and sub-pallium. At embryonic day 15.5, Slc38a1 localizes in neuronal stem cells in the ventricular zone in the forebrain, while it localizes in mature neurons in the sub-ventricular zone of the hindbrain. In the adult brain, Slc38a1 is not detected in neuronal stem cells in the sub-granular zone, however, it is highly enriched in mature local parvalbumin^+^ and somatostatin^+^ interneurons and ependyma. Analyses of single-cell RNA sequencing data further reveal that both the number of Slc38a1-expressing cells and the average transcript level of Slc38a1 are significantly higher in embryo brain cells compared to adults. In the embryo, Slc38a1 transcripts are particularly enriched in immature neuronal lineages and embryonic oligodendrocyte precursor populations, whereas in the adult, expression is more prominent in neural progenitor cells and radial glia. Disruption of *Slc38a1* in mice impacts body weight, brain size and glutamatergic neuronal volume, while dendritic arborization is diminished and cellular life span is shortened. Altogether, Slc38a1 is essential for normal neurodevelopment, indirectly regulates adult neurogenesis in the sub-granular zone and influence neuronal morphology and cell viability.

## INTRODUCTION

The adult human brain consists of ∼86 billion neurons (1), necessitating the generation of ∼250,000 neurons per minute in the embryonic human brain (2). Projection neurons are generated in the cortical preplate of the dorsal pallium and migrate radially to the cortical plate (3). In contrast, interneurons originate in the sub-pallium and migrate tangentially to integrate with projection neurons in the cortical plate (4). In the adult brain, neurogenesis persists in restricted areas, including the sub-granular zone (SGZ) of the hippocampus — which contributes to granule cell replacement and new circuit formation, the sub-ventricular zone (SVZ) of the lateral ventricle wall —where neurons migrate along the rostral migratory stream to the olfactory bulb—and the hypothalamus, where it supports metabolic and circadian regulation (5, 6). Neurogenesis depends on ultrafast supplies of energy, amino acids and building blocks such as proteins and nucleotides. However, how the neurogenic niche is furnished with these components and how neurogenesis is regulated in both embryonic and adult contexts, remains largely unresolved. Emerging evidence highlights a critical role for the amino acid glutamine in neurogenesis. Glutamine enhances self-renewal in mouse embryonic stem cells (ESCs) and helps maintain their undifferentiated state (7). Glutaminolysis promotes basal progenitor generation and contributes to cortical plate expansion and gyrification in both humans and mice (8, 9). The glutamine-derived neurotransmitters glutamate and GABA also regulate early neurodevelopment (10). Despite the fundamental functions of these amino acids in the embryonic brain, the mechanisms involved in glutamine supply and its impact on the embryonic and adult neurogenic cells have eluded characterization.

The Solute carrier family 38 (Slc38) consists of 11 homologous neutral amino acid transporters translocating neutral amino acids, including glutamine, across (sub)cellular membranes (11, 12). Among these, Slc38a1 (also known as (aka) SAT1, SNAT1, SA2, and ATA2) is remarkable by having high affinity for glutamine (Km=0.37 mM at −50 mV in *Xenopus laevis* oocytes) and ability to accumulate high levels of glutamine intracellularly (13, 14). In the adult, Slc38a1 supplies parvalbumin^+^ (PV^+^) interneurons with glutamine for GABA synthesis and impacts vesicular morphology and GABA load (15, 16). Genetic inactivation of Slc38a1 in mice leads to impaired critical period plasticity, changed high-frequency membrane oscillations and perturbed cortical processing and plasticity in the adult (15, 16).

Interestingly, Slc38a1 is upregulated in various cancers and has been associated with poor prognosis (17, 18). This aligns with the “glutamine addiction” hypothesis (19), which posits that rapidly dividing cancer cells depend heavily on glutamine to meet their increased demands for energy, amino acids, nucleotides, and other biosynthetic precursors. Given the parallels between cancer cells and neurogenic cells—particularly their rapid proliferation and reliance on glutamine-derived substrates for metabolism and growth, we hypothesized that Slc38a1 may play a key role in sustaining neurogenesis. Here, we show that Slc38a1 transcript and protein are widely expressed in the embryonic brain, including both the dorsal pallium and subpallium. This suggests critical roles for Slc38a1 in the embryonic neurodevelopment of glutamatergic projection neurons and GABAergic interneurons. In the adult brain, neural stem cells within the SGZ do not express Slc38a1; however, the transporter is highly enriched in local PV^+^- and somatostatin^+^ (SST^+^)-, indicating an indirect role in regulating adult neurogenesis. Genetic inactivation of Slc38a1 leads to significant phenotypic alterations: reduced body weight and brain length, decreased packing density of hippocampal pyramidal and granule cells, simplified neuronal morphology and a shortened cellular life span.

## MATERIALS AND METHODS

Detailed description of all methods, materials and procedures are provided in the Supplementary Information.

### Animals

All animal procedures were carried out in full compliance with the Norwegian Animal Welfare act and the European Union Directive 2010/63/EU on the protection of animals used for scientific purposes. The study was approved by the Norwegian Food Safety Authority (FOTS application numbers 9200, 21009 and 30120) and reported in accordance with the ARRIVE guidelines for reporting animal research (20). Mice were housed under controlled conditions with a 12-hour light/dark cycle, ambient temperature maintained at 22 ± 1 °C, and relative humidity at 50 ± 10%. Animals had ad libitum access to autoclaved water and RM3 chow (Special Diets Service, UK). All transgenic mice used in the study—both male and female—had been backcrossed onto the C57BL/6J background for at least 10 generations.

The *Slc38a1*^−/−^ mouse model used in these studies has previously been rigorously characterized phenotypically (15,16). Briefly, the floxed *Slc38a1* targeting construct was generated by recombination technique. A 12.9 kb genomic region containing exons 5–12 was isolated from BAC clone RP23-85D13, and LoxP sites were inserted into introns 4 and 8 along with an FRT-flanked PGKNeo cassette in intron 8. This design floxed a 4.5 kb region spanning exons 5–8, such that Cre recombination causes a frameshift deletion. The targeting construct was electroporated into D1 ES cells (129S6 × C57BL/6J), and G418-resistant clones were identified by nested PCR using primers outside the homology arms and within the neo cassette. Chimeric mice were produced by aggregating targeted ES cells with CD-1 eight-cell embryos. The neo cassette was excised by crossing chimeras with homozygous FLP-expressing 129S4/SvJaeSor-Gt(ROSA)26Sortm1(FLP1)Dym/J females. F1 offspring were genotyped by PCR using primers flanking the LoxP or FRT-LoxP sites. These mice were subsequently crossed with Deleter mice (21) to generate a constitutive *Slc38a1* knockout.

### Quantitative western blotting

Quantitative western blotting was performed as previously described (18, 21) with minor modifications. Briefly, brains were collected from three age- and sex-matched pairs of Slc38a1^+/+^ and Slc38a1^−/−^ mice and rats at defined developmental stages. Animals were anesthetized and decapitated; brains were rapidly dissected, snap-frozen on dry ice, and stored at –80 °C until further processing. Brain tissue was homogenized and protein extracts were separated by SDS-PAGE and transferred onto PVDF membranes. Membranes were incubated overnight at 4 °C with primary antibodies against Slc38a1 (home-made; (22)) or GAD67 in 3% skim milk in Tris-buffered saline with 0.1% Tween-20 (TBST). Blots were imaged using a ChemiDoc Imaging System (Bio-Rad), and band intensities were quantified following background subtraction. Signal intensities of Slc38a1 bands were normalized to GAPDH on the same blot. Data from all age groups were further normalized to the mean normalized signal of the adult group to allow cross-age comparison.

### Immunofluorescence staining

Three age- and sex-matched pairs of Slc38a1^+/+^ and Slc38a1^−/−^ mice, along with three rats at defined developmental stages, were anaesthetized and transcardially perfused with freshly prepared 4% paraformaldehyde (PFA) in phosphate-buffered saline (PBS). Embryonic mice were not perfused but instead immersion-fixed. Brains were rapidly dissected, and the size of the left or right hemisphere was measured using a scaled ruler on paper. Brains were then immersion-fixed in 4% PFA overnight at 4 °C. After fixation, tissues were cryoprotected by sequential incubation in 10%, 20%, and 30% sucrose solutions in PBS until fully submerged. Brains were then embedded in Neg-50™ Frozen Section Medium and cryosectioned at 18 μm thickness in either the coronal or sagittal plane.

Sections were incubated overnight at 4 °C with one or two primary antibodies of interest, followed by washing and incubation with appropriate species-specific fluorescent secondary antibodies. Nuclei were counterstained with DAPI, and sections were mounted using ProLong™ Gold Antifade Mountant. Images were acquired and analyzed using a Zeiss LSM700 confocal microscope system.

### RNA detection (*in situ* hybridization) with RNAscope probe in embryonic and adult mouse brain

Brain sections from age- and sex-matched Slc38a1^+/+^ and Slc38a1^−/−^ mice were prepared as for immunofluorescence staining. Localization of mRNA was performed using the RNAscope® 2.5 High Definition (HD)-BROWN Assay kit, according to the manufacturer’s instructions. Hybridization signals were detected using 3,3′-diaminobenzidine. Slices were mounted with Eukitt mounting medium and analyzed with a Leica microscope. The following RNAscope probe was used for detection of Slc38a1 mRNA: RNAscope® Probe - Mm-Slc38a1-C1, Mus musculus solute carrier family 38 member 1 (Slc38a1) transcript variant 1 mRNA. In addition, RNAscope Negative Control Probe-DapB, and RNAscope® Positive Control Probe – PPIB were used.

### Data curation and analysis

Single-cell RNA sequencing datasets representing mouse brain nuclei from adult (23) and embryonic (24) stages were obtained from Parse Biosciences via the Trailmaker™ dataset repository (25). The datasets were provided as preprocessed .rds files, accompanied by corresponding metadata. Quality control and downstream analysis were performed using Scanpy (26) (v1.9.8). Cells expressing fewer than 200 genes and genes expressed in fewer than 15 cells were excluded. Additional filtering involved the removal of outlier cells based on “log1p_total_counts”, “log1p_n_genes_by_counts”, and “pct_counts_in_top_20_genes”, using thresholds of 8 median absolute deviations (MADs) for the adult dataset and 6 MADs for the embryonic dataset. Normalization and identification of highly variable genes (n_top_genes set to 2000) were performed using Sanpy’s built-in functions. The final dataset included 7361 cells from the adult mouse brain and 15954 from the embryonic brain. Cluster annotations were obtained directly from the accompanying metadata.

### Hippocampal granule cell layer (GCL) and CA1 pyramidal cell layer (PCL) width measurement

A series of sagittal 18 μm thick sections were made by a cryostat exhibiting structures of both upper GCL and PCL (27). To measure the width of GCL and PCL of hippocampus, their border was defined as the border of the last nuclear contact to the main layer. Seven randomized and anatomically comparable regions of interest were selected from the upper DG and CA1 region of the hippocampus for quantification. Measurements of layer thickness were performed using Adobe Photoshop CS6, utilizing the measurement tool to ensure precision and consistency across sections.

### Neuron survival counting

Cortical tissue from embryonic day 18.5 (E18.5) Slc38a1^+/+^ and Slc38a1^−/−^ mice was carefully dissected, dissociated, and cells were placed into a dish with complete medium (Neurobasal + B27). The primary cortical cells were incubated at 37°C and 5% CO_2_ and the cell medium changed every third day. The neuronal cultures matured after 2 weeks’ incubation.

At the appropriate time point, primary neurons were fixed and incubated overnight at 4 °C with an antibody against MAP2 to label dendritic structures. For quantification, seven randomized confocal microscope views (20x) of each cover glass were captured. Cells within each field were manually counted in a blinded manner using the counting tool in Adobe Photoshop CS6. Cell counts at DIV2 and DIV14 were normalized to the average counts of Slc38a1^+/+^ neurons of at the corresponding time point.

### Sholl analysis of dendritic complexity

Sholl analysis was conducted using the Simple Neurite Tracer plugin in Fiji/ImageJ, following the guidelines provided in the ImageJ user manual (imagej.net). This semi-automated tool enables tracing, visualization, and quantitative analysis of neuronal processes.

Neuronal dendrites were first traced manually using the plugin, starting from a defined origin and increasing in radius by 10 μm increments, were then overlaid on the traced images. The starting radius was placed at the origin of the longest dendrite for each individual neuron. Intersections were defined as the points where neuronal processes crossed the concentric circles.

A total of 21 MAP2⁺ neurons per genotype were randomly selected from 14-day in vitro (DIV14) cultures and imaged at 40× magnification. For each neuron, the number of intersections, dendritic branch points, and total dendritic length were quantified, following protocols described in previous studies (28).

### Statistical analysis

All data are presented as mean ± SEM. Statistical analyses were performed using GraphPad Prism 9. Depending on the experimental design, either non-parametric *t*-test or Two-way ANOVA were applied as appropriate. A *p*-value of < 0.05 was considered statistically significant.

## RESULTS

### Slc38a1 protein expression in the germinal zones of the embryonic rat brain

To investigate the role of Slc38a1 in the developing brain – particularly at a stage where neuronal proliferation is at a peak and requires high levels of nucleotides, amino acids, signaling molecules and more, we first compared Slc38a1 protein expression in embryonic and adult rat brains. Using our highly specific anti-Slc38a1 antibody (22), Western blot analysis revealed a single band at ∼50 kDa in both embryonic and adult brains (Fig. 1A; S1A), consistent with previous reports (15). Quantification indicated that the expression of Slc38a1 protein levels at embryonic day 15.5 (E15.5) were comparable to those observed in the adult brain (Fig. 1B). In contrast, GAD67, a key enzyme in the formation of neurotransmitter GABA, was barely detected in the embryonic brain (Fig. 1C-D). Given that classical synaptic transmission is not yet established at E15.5, while neurogenesis is at its peak (29), these findings suggest that Slc38a1 may serve a distinct, developmentally relevant function during embryonic neurogenesis.

**Figure 1.**
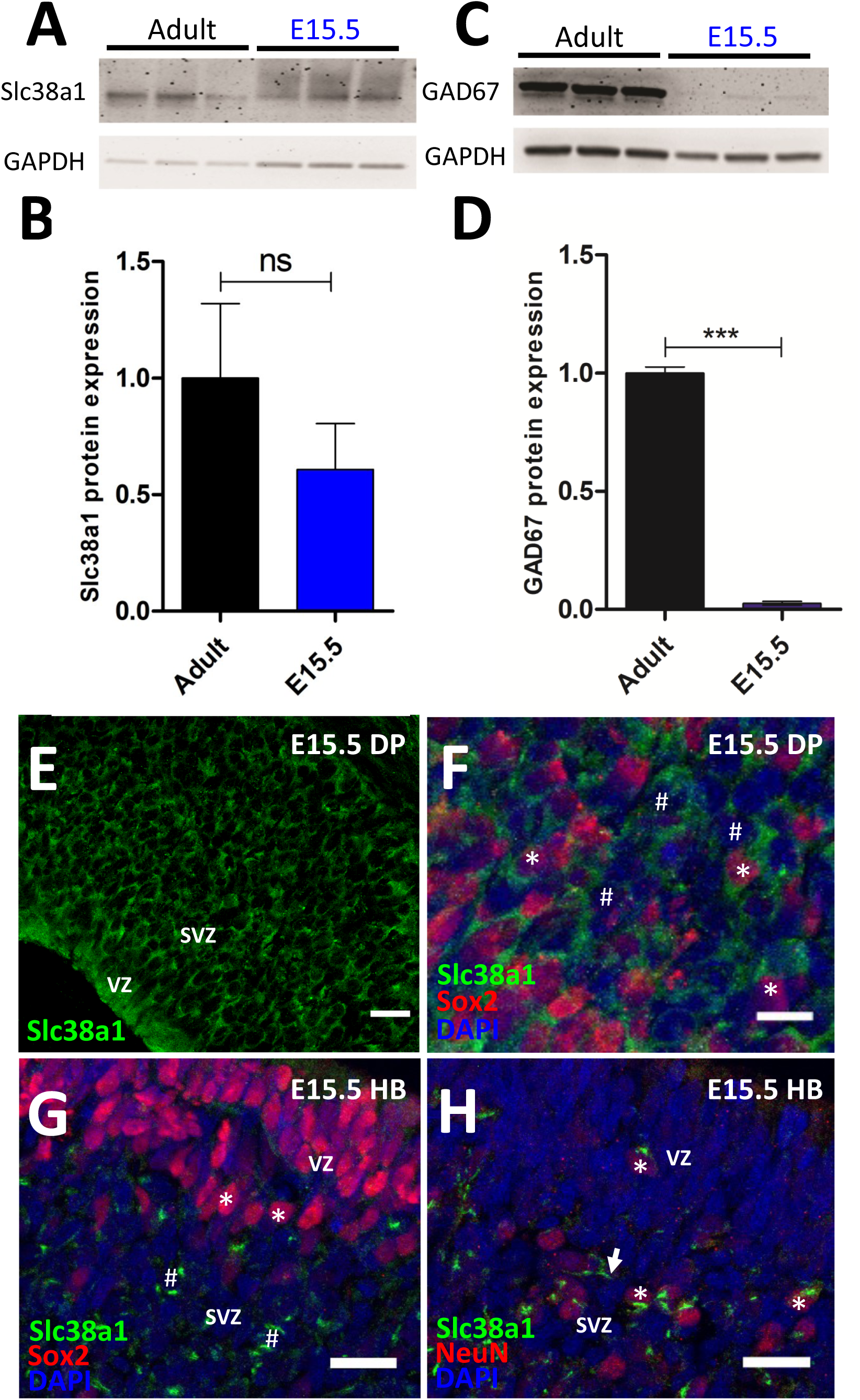
Slc38a1 protein expression in the germinal layers of embryonic rat brain – localized in stem/progenitor cells and in mature neurons at E15.5. (A-D) Three whole rat brain extracts from the embryonic day 15.5 (E15.5) and three extracts from adult were electrophoretic separated in SDS-PAGE followed by immuno-labeling for Slc38a1 or glutamate decarboxylase 67 (GAD67). Slc38a1- and GAD67-labeling was quantified by comparing to the house-keeping gene glyceraldehyde-3-phosphate dehydrogenase (GAPDH) followed by normalization to the mean value of the adults. (A) Anti-Slc38a1 antibody reveals a specific band at ∼50 kDa, both in the embryonic and in the adult brains. (B) Slc38a1 protein expression levels in the rat embryonic brain and the adult rat brain show no significant difference. (C) Anti-GAD67 antibody shows a powerful band at ∼67 kDa in the adult brain, while there is barely any band of the same size in the embryonic brain tissue. (D) GAD67 protein expression in the E15.5 rat brain is almost non-existing compared to expression in the adult rat brain. (E-H) E15.5 embryonic rat brains were triple-labeled for Slc38a1 (green), cell-specific markers (red) and DAPI (blue) and investigated by confocal laser scanning microscopy. (E) Slc38a1 staining appears as cell-like structures in the ventricular zone (VZ) and the sub-ventricular zone (SVZ) of the rat embryonic dorsal pallium (DP). (F) Merged and enlarged images of Slc38a1, Sox2 and DAPI staining in DP show that Slc38a1 labeling surrounds Sox2^+^/DAPI^+^ (*) as well as Sox2^−^/DAPI^+^ (^#^) stained cell nuclei. (G) In the hindbrain (HB), Slc38a1 labeling is mostly in cytoplasm around Sox2^−^/DAPI^+^ nuclei (^#^) in the SVZ, while the cytoplasm of Sox2^+^/DAPI^+^-cells (*) in the VZ is devoid of Slc38a1 staining. (H) In most occasions in the HB, Slc38a1 staining is enriched asymmetrically around NeuN^+^/DAPI^+^-nuclei (*) and extends into neuronal processes (arrow) in the SVZ. Sox2 is a neuroepithelial stem cell and progenitor cell marker, and NeuN (aka Sox3) is a marker for mature neurons. (Note that Sox2 is a transcription factor and NeuN a splicing regulator, showing nuclear staining.) DAPI visualizes nuclear DNA. Scale bars represent 20 μm (F, G, H) and 10 μm (E). Data are presented as mean+SEM. ***, P=0.001, the unpaired *t*-test statistics was done by Prism GraphPad 9.

To determine the cellular localization of Slc38a1 in the embryonic rat brain we performed immunohistochemistry on E15.5 rat brain sections. Robust, cell-like Slc38a1 staining, suggesting the localization of Slc38a1 in the cytoplasm of cells and in the cell membranes, was detected in both ventricular zone (VZ) and SVZ of the dorsal pallium (Fig. 1E) – the region where cortical glutamatergic neurons and glial cells are born (30). Co-staining with Sox2, a marker for multipotential neuronal stem cells (NSCs) and progenitor cells (31), revealed overlapping expression in the same regions. Merged images of Slc38a1, Sox2 and the nuclear stain DAPI showed Slc38a1 localization surrounding both Sox2^+^/DAPI^+^ and Sox2^−^/DAPI^+^ nuclei (Fig. 1F; S1B), suggesting that Slc38a1 is expressed in neural stem cells, progenitor and post-mitotic cells. Similar expression patterns were observed in the VZ and SVZ of the lateral ganglionic eminence (LGE), and the medial ganglionic eminence (MGE) – two brain regions giving birth to a range of GABAergic interneurons (Fig. S1C-D; (32)). In contrast, the embryonic hindbrain stands as an opposite: Slc38a1 staining segregated to Sox2^−^/DAPI^+^-cells in the SVZ, while DAPI^+^/Sox2^+^ cells in the VZ were devoid of Slc38a1 staining (Fig. 1G). Supporting this, Slc38a1 staining did not co-localize with the neuroepithelial stem cell protein Nestin in the hindbrain (Fig. S1E), but instead was localized peri-nuclear and along the processes of NeuN^+^ mature neurons cells (Fig. 1H). Collectively, these findings demonstrate that Slc38a1 is enriched in germinal zones of the embryonic brain, particulary in regions responsible for producing glutamatergic, GABAergic and glial cells lineages. Moreover, while Slc38a1 is present in immature NSCs of the VZ and SVZ in the forebrain, it is confined to mature SVZ neurons in the hindbrain at E15.5, consistent with the earlier maturation of hindbrain structures compared to the forebrain (33).

### Robust peri-natal Slc38a1 staining in the mouse brain

To further support our observations of embryonic Slc38a1 localization and its potential role in brain development, we examined Slc38a1 protein levels across six developmental stages − from the embryonic to the adult − in Slc38a1^+/+^ and in Slc38a1^−/−^ mice using Western blotting. A single ∼50 kDa band was detected in all six groups of Slc38a1^+/+^ mice (Fig. 2A). Quantitative immunoblotting confirmed that Slc38a1 protein levels at E15.5 and adult mouse brain are similar (Fig. 2B), in harmony with findings in the rat brain (Fig. 1A-B). Interestingly, Slc38a1 levels increase significantly from E15.5 to peak at P7, when synaptogenesis is highest (34), and then decrease steeply to reach the adult levels at P20. No stained bands were detected on Slc38a1^−/−^ mice at any stage (Fig. 2A-B), verifying successful inactivation of Slc38a1 and the specificity of the Slc38a1 antibody.

**Figure 2.**
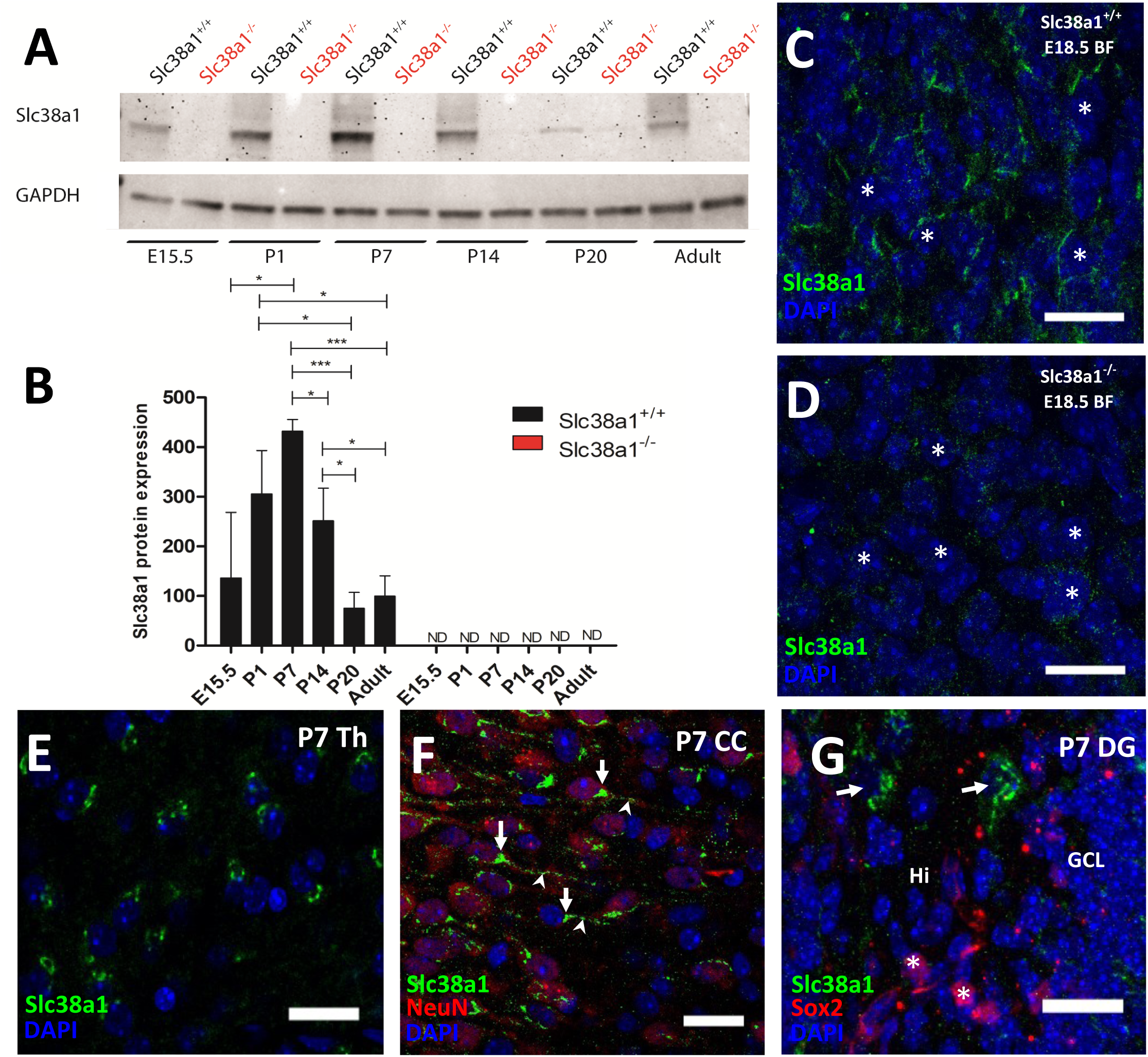
Robust Slc38a1 protein expression in the prenatal (E) and postnatal (P) mouse brain peaking at P7. (A-B) Slc38a1 protein levels were investigated in whole brain samples from six developmental stages in mice by western blotting. (A) In Slc38a1^+/+^ mice, immunoblots show staining for Slc38a1 in all six stages, from the embryonic day 15.5 (E15.5) to adult mice, while no expression is detected in the same six stages in Slc38a1^−/−^ mice. (B) Slc38a1 protein levels were quantified by comparing Slc38a1 signal intensity to the house-keeping gene GAPDH and normalized to mean value of adult mice. In Slc38a1^+/+^ mice brains, Slc38a1 expression increases significantly from E15.5 to a peak at P7, and then decreases to the same levels as E15.5 in the adult mouse brain. No expression of Slc38a1 is detected in any of these stages in Slc38a1^−/−^ mice. The bar graph shows mean+SEM. *, P=0.05, ***, P=0.001. The unpaired *t*-test statistics was done by Prism GraphPad 9. ND: not detected. (C-G) Sections from embryonic day 18.5 (E18.5) and P7 brain were triple stained for Slc38a1 (green), cell-specific markers (red) and DAPI (blue) and analyzed by confocal laser scanning microscopy. (C-D) Antibody against Slc38a1 labels cells around DAPI^+^-nuclei (*) in the embryonic basal forebrain (BF) of Slc38a1^+/+^ mouse. The same antibody does not visualize labeled structures around DAPI^+^-nuclei (*) in the BF of embryonic Slc38a1^−/−^ mouse brain, except for some background. (E) Rosette-like staining for Slc38a1 surrounding DAPI^+^-nuclei is detected in the thalamus (Th) at P7. (F) In the mouse cerebral cortex (CC) at P7, staining for Slc38a1 localizes in NeuN^+^ neurons. The Slc38a1 staining is outside the nuclei, often concentrated at a pole (arrow) and extending discontinuously into neuronal processes (arrowhead). (G) In the P7 dentate gyrus (DG), Slc38a1 labeled cells are concentrated in the hilus (arrow). The staining does not associate with Sox2^+^ cells (*). No staining for Slc38a1 is detected in the granule cells of the DG. The Slc38a1 staining (C-G) is compatible with localization mostly in cell membranes and cytoplasmic organelles in neurons. Sox2 is a neuroepithelial stem cell and progenitor cell marker, while NeuN is a marker for mature neurons. Scale bars represent 20 μm (C-G). Hi: Hilus of the DG; GCL: granule cell layer.

We next explored the cellular staining of Slc38a1 during the peri-natal period using immunofluorescence. Slc38a1-like staining is detected in basal forebrain of Slc38a1^+/+^ mice in the late embryonic stage (Fig. 2C), while such staining was not seen in the same region of Slc38a1^−/−^ mice of the same age (Fig. 2D), confirming specific Slc38a1 staining in the wild type mice. In the early postnatal brain stem, Slc38a1 labeling forms rosette-like structures around nuclei in thalamus of Slc38a1^+/+^ mice (Fig. 2E). In the cerebral cortex of P7 mice, NeuN^+^ cells are decorated with peri-nuclear cup-like staining for Slc38a1, which elongates into neuronal processes (Fig. 2F). Slc38a1 antibody stains scattered cells in the hilus of the DG, but this staining does not colocalize with Sox2^+^ nuclei (Fig. 2G). Altogether, these data show widespread, but select, perinatal Slc38a1 protein staining in wild-type mice, in harmony with findings in the rats.

### In the adult brain, Slc38a1 protein is absent from nestin^+^/GFAP^+^ sub-granular cells in the DG, but expressed in ependymal cells and interneurons

Given the strong localization of Slc38a1 in embryonic germinal zones, we next asked whether it might also play a role in adult neurogenesis. We focused on the DG of the adult hippocampal, a well-established neurogenic niche (30). In this region, there are multiple Slc38a1^+^, Nestin^+^ and GFAP^+^ cells, but no co-localization of Slc38a1 with GFAP and Nestin is detected (Fig. 3A, S2A). Slc38a1 is localized in most ependymal cells in the wall of the lateral ventricle (LV) of adult Slc38a1^+/+^ mice (Fig. 3B, S2B). A subset of Slc38a1^+^ ependymal cells co-localize with Sox2 in the ependymal layer, but not in SVZ (Fig. 3C). The specificity of the Slc38a1 staining is supported by deprived Slc38a1 staining of ependymal cells in the adult Slc38a1^−/−^ mice (Fig. S2C).

**Figure 3.**
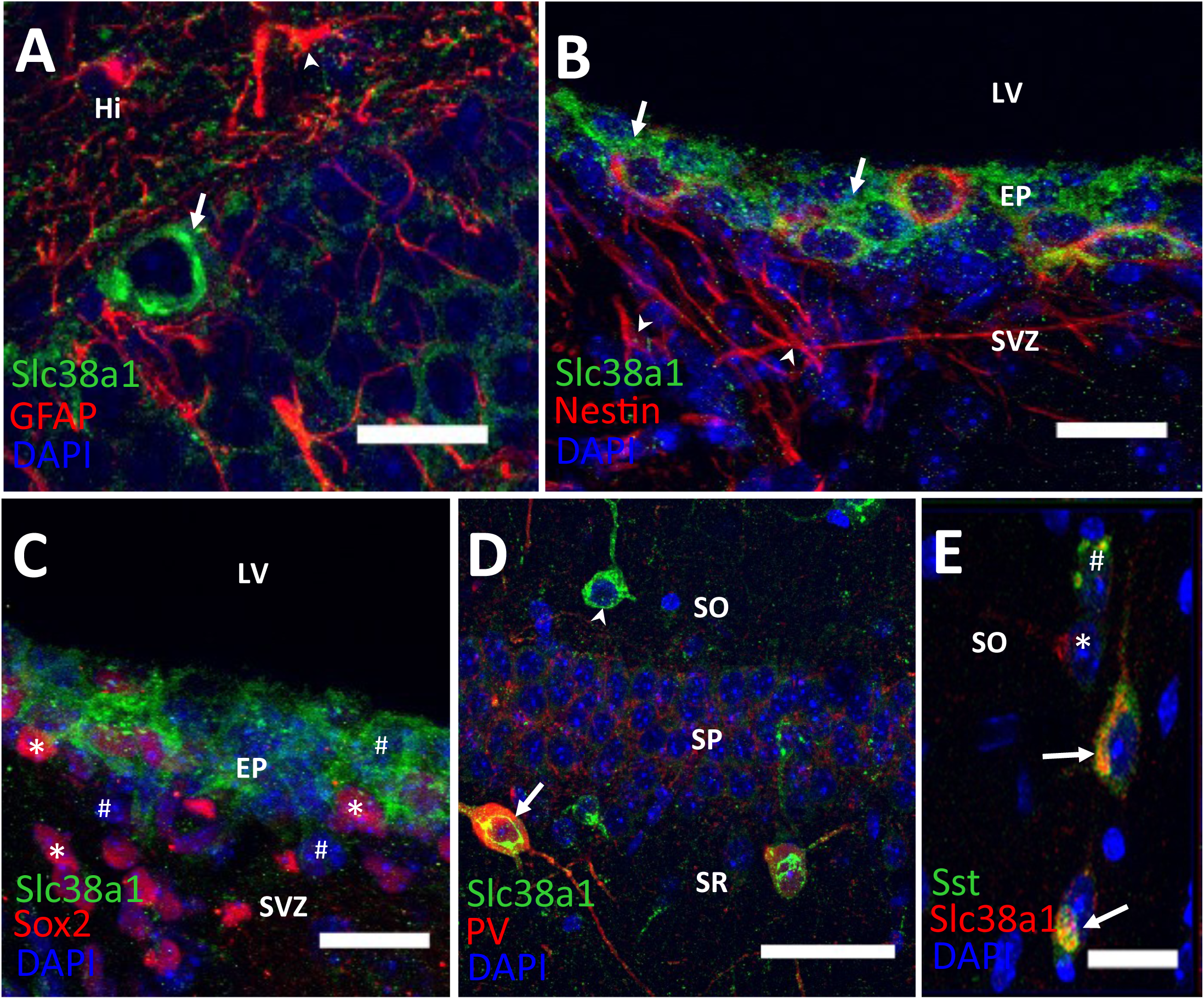
Slc38a1 protein expression appears in adult parvalbumin^+^ (PV), somatostatin^+^ (SST) and ependymal cells, but not in dentate gyrus (DG) progenitor cells. (A-E) Brain tissue from adult mice were labeled for Slc38a1 (green/red), cell-specific markers (red/green) and DAPI (blue) and investigated for a role of Slc38a1 in adult neurogenesis. (A) Slc38a1 staining (arrow) shows strong labelling in the cytoplasm and cell membrane in scattered cells (likely basket interneurons) and much weaker labelling of granule cells in the DG of the hippocampal formation in Slc38a1^+/+^ mice. It does not co-localize with GFAP^+^ cells, i.e. astrocytes (arrowhead). (B) Slc38a1 protein is expressed in most ependymal cells (arrows) lining the lateral ventricle (LV) of Slc38a1^+/+^ mice, but co-localizes minimally with nestin^+^ ependymal cells (red). No staining for Slc38a1 is detected in the sub-ventricular zone (SVZ). (C) Slc38a1 co-localizes with Sox2^+^ cells (*) and Sox2^−^ cells (^#^) in the ependymal layer in adult Slc38a1^+/+^ mice. No staining for Slc38a1 is detected in Sox2^+^ cells or Sox2^−^cells in the SVZ. (D) In the hippocampal CA1, Slc38a1 staining is seen in scattered neuron-like cells in the oriens (SO), pyramidal (SP) and radiatum (SR) layers. Some Slc38a1-labeled cells in or around SP co-express PV (arrow), while Slc38a1 labeled cells in the SO do not co-express PV (arrowhead). (E) In the CA1, some interneurons express Slc38a1 (*), some SST (^#^) and some both Slc38a1 and SST (arrow). Scale bars represent 20 μm.

Anti-Slc38a1 antibody also stains scattered cells in the adult mouse CA1 (Fig. 3D; S2D), consistent with our previous publication (15). Surprisingly, many Slc38a1^+^ cells in the stratum oriens do not co-stain for PV (Fig. 3D). As SST^+^ subgroup of interneurons is particularly enriched in the stratum oriens and hilus of the DG (35), we investigated whether Slc38a1^+^ cells contained SST. Indeed, Slc38^+^ cells co-stain PV, SST or none (Fig. 3E). Thus, we demonstrate that Slc38a1 is not limited to PV^+^ interneurons in the adult.

Slc38a1 resides in endomembrane system, with a major fraction in post Golgi secretory pathway (Fig. S3A, S3C). Its predominant localization in Trans-Golgi, plasma membrane and absence in cis Golgi suggest that it shuttle between plasma membrane and Golgi (Fig. S3A, S3C-D). We also observed Slc38a1 in endoplasmic reticulum (ER), gateway to endomembrane system and late endosomal compartment (CD63) as well as early endosomes (EEA1) (Fig. S3B, S3E-F).

### Ample Slc38a1 transcript in embryonic mouse brain supports a role for Slc38a1 in neurogenesis

To further rule out spurious data due to, e.g., possibly unspecific antibody or errors in the immunocytochemistry experiments, we reinforced our findings by investigating Slc38a1 transcript localization by *in situ* hybridization. Complementary RNA probes (anti-sense) applied on coronal sections from E18.5 mice show high level of Slc38a1 transcript in the dorsal pallium and cerebral cortex of Slc38a1^+/+^ mice (Fig. 4A, C; S4A), consistent with results obtained for Slc38a1 protein by immunofluorescence assays (Fig. 1E-F). On the contrary, significantly lower transcript signal was found in the dorsal pallium and cerebral cortex from Slc38a1^−/−^ mice (Fig. 4A-C). This confirms the specificity of the probe used to detect the mRNA of Slc38a1. High Slc38a1 transcript levels were detected in many brain regions, including the LGE, choroid plexus, the mammillary nucleus and lateral hypothalamus (Fig. 4D, S4B-C). In summary, *in situ* hybridization in the embryonic brain provides additional evidence for ample expression of Slc38a1 in various parts of the developing brain bolstering its role in brain development.

**Figure 4.**
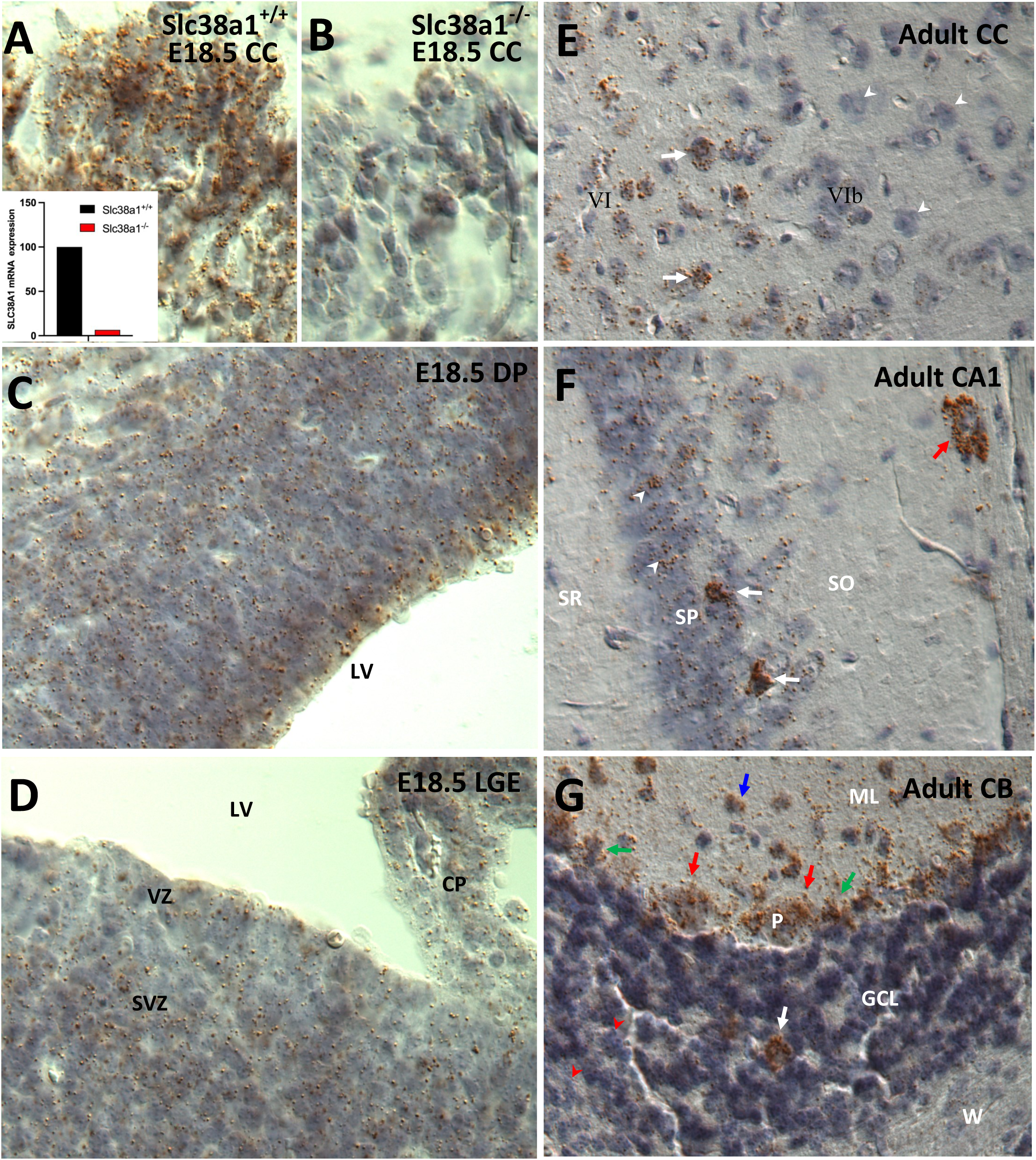
Slc38a1 transcript is widely distributed in the embryonic mouse brain, while in the adult brain it is concentrated in select cells. Coronal sections were made of E18.5 and adult mouse forebrain, labeled with complementary RNA (anti-sense) for Slc38a1 (*in situ* hybridization) and the localization investigated. (A, B) Strong labeling (brown precipitate) for Slc38a1 mRNA is detected in the cerebral cortex (CC) of Slc38a1^+/+^ mice. Few probe particles are seen in the corresponding part of CC in Slc38a1^−/−^ mice. Inset in A shows quantification of probe labeling in A compared to B. (C) Slc38a1 antisense labels throughout the dorsal pallium (DP). (D) the lateral ganglion eminence (LGE) is strongly decorated with Slc38a1 probe particles, the highest concentration being in the sub-ventricular zone (SVZ). Choroid plexus (CP) is also labeled for Slc38a1 transcript. (E) In the adult CC, the Slc38a1 probe labels distinct cells. A large number of cells in the layer VI show strong Slc38a1 transcript (arrows), while most cells in VIb are devoid of labeling (arrowhead). (F) In the adult hippocampal CA1, Slc38a1 probe labeling is strongly enriched in some scattered cells in the pyramidal layer (SP; white arrow) and at the external border of the oriens layer (SO; red arrow). Lower labeling is detected on pyramidal cells (arrowhead). (G) In the adult cerebellum (CB), Purkinje cells (P; red arrows) and some scattered cells in the granule cell layer (GCL; white arrow) and molecular layer (ML; blue arrow) are strongly labeled by Slc38a1 probes. Small, stained cells between Purkinje cells are Bergman glia (green arrow). Most cells in the granule cell layer show less labeling for Slc38a1 transcript (red arrowhead). SR: stratum radiatum; W: white matter.

### Slc38a1 mRNA is enriched in distinct, matured local interneurons and ependymal cells in the adult mouse

In the adult mouse brain, *in situ* hybridization revealed strong Slc38a1 transcript labeling in layer VI of the cerebral cortex (Fig. 4E), a region where >75% of neurons are PV^+^ and/or SST^+^ (36). In contrast, the Slc38a1 mRNA was sparse in layer VIb – a region with low PV^+^- and SST^+^-interneurons (Fig. 4E). In the hippocampus, Slc38a1 mRNA was strongly expressed in scattered cells located in the stratum pyramidale, stratum oriens and in the dentate hilus (Fig. 4F, S4D). Similarly, in the cerebellum, Slc38a1 transcripts was enriched in Purkinje cells, Bergmann glial cells and some scattered cells in the molecular and granular layers (Fig. 4G). These scattered cells likely correspond to stellate and basket cell in the molecular layer, which are GABAergic, and Golgi cells in the granular layer, which are both GABAergic and glycinergic (37). Medium level Slc38a1 transcript is detected in the hippocampal pyramidal and granule cells and cerebellar granule cells (Fig. 4F, G; Fig. S4D), which are all glutamatergic. Consistent with our demonstration of Slc38a1 protein expression in ependymal cells, *in situ* hybridization confirms Slc38a1 transcript localized in ependymal cells in the adult, while cells in the choroid plexus do not reveal Slc38a1 transcript (Fig. S4E, F). Thus, the *in situ* hybridization localization of Slc38a1 in the adult rat brain mirrors for the most part the Slc38a1 protein distribution (15, 22).

### Single-cell RNA sequencing supports a role of Slc38a1 in neurogenesis

To further support our findings implicating Slc38a1 in neurogenesis and to unravel Slc38a1 expression dynamics in individual cells and in time, we curated and analyzed publicly available datasets from Parse Biosciences (23–25). Data were visualized using Uniform Manifold Approximation and Projection (UMAP), separated into embryonic and adult brain samples (Fig. 5A-B). Feature plots of Slc38a1 transcript levels revealed higher expression and broader distribution in the embryonic brain compared to the adult brain (Fig. 5C–E). In embryoni c tissue, Slc38a1 expression was most prominent in neural and glial progenitor populations, including immature neurons and oligodendrocyte precursor cells (OPCs) (Fig. 5F). In the adult brain, Slc38a1 was also expressed in progenitor populations, such as neuronal progenitor cells, radial glial cells, and neuroepithelial cells (Fig. 5G).

**Figure 5.**
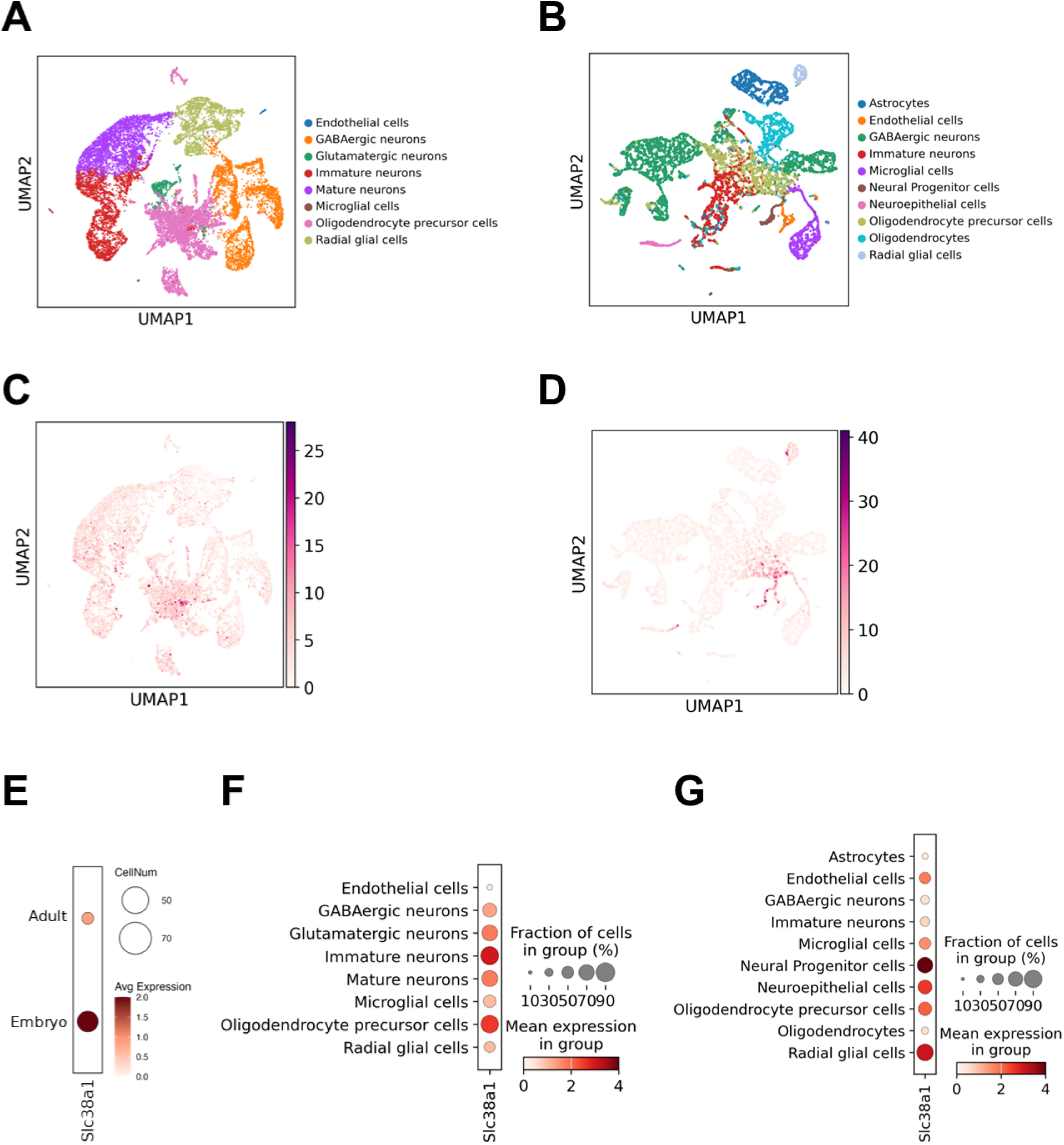
Single cell RNA sequencing demonstrates strong Slc38a1 transcript in multiple embryonic and adult cells. Single-cell data from brain embryos and adults, obtained from publicly available Parse Biosciences datasets (23–25) were analyzed. UMAP plots display the cell clusters in embryonic (A) and adult (B) mice brain samples. Feature plots of Slc38a1 gene expression highlight its distribution in embryonic (C) and adult (D) brain cells, while a dot plot summarizes the average expression levels across identified cell types in both developmental stages (E–G). Overall, both the number of Slc38a1-expressing cells and the average transcript levels were higher in the embryonic brain compared to the adult brain (E). In embryos, Slc38a1 transcript was most enriched in immature neurons and oligodendrocyte precursor clusters (F). In adults, the highest expression was observed in neural progenitor, radial glial and neuroepithelial cells (G). Comparative analysis between embryonic and adult brains revealed that immature neurons, glutamatergic and GABAergic neurons and oligodendrocyte precursor cells showed higher Slc38a1 RNA expression in the embryonic brain (F), whereas radial glial cells displayed relatively higher expression in the adult brain (G).

Comparative analysis revealed that Slc38a1 expression is significantly upregulated in embryonic populations, particularly in immature neurons, oligodendrocytes, and both glutamatergic and GABAergic neurons, consistent with a developmental role in sustaining neurogenesis. As expected, Slc38a1 expression was negligible in non-neuroectodermal cell types, such as endothelial cells and microglia, which are derived from the mesoderm and do not participate in neurogenesis. In summary, our scRNA-seq analysis highlights the dynamic and cell-specific expression of Slc38a1, reinforcing its critical role in both embryonic and adult neurogenic processes.

### Slc38a1 disruption impacts body weight, brain size and neuronal number

To assess the physiological consequences of Slc38a1 disruption, we monitored body weight in male mice at three different ages. At 10 weeks of age, no significant difference in body weight was observed between Slc38a1^+/+^ and Slc38a1^−/−^ mice males. However, starting at 13 weeks, Slc38a1^−/−^ displayed a significant reduction in body weight compared to wild-type controls, a difference that became more pronounced by 30 weeks of age (Fig. 6A). Similarly, female Slc38a1^−/−^ mice at 13 weeks exhibited significantly lower body weight compared to their *Slc38a1*^+/+^ counterparts (Fig. 6B).

**Figure 6.**
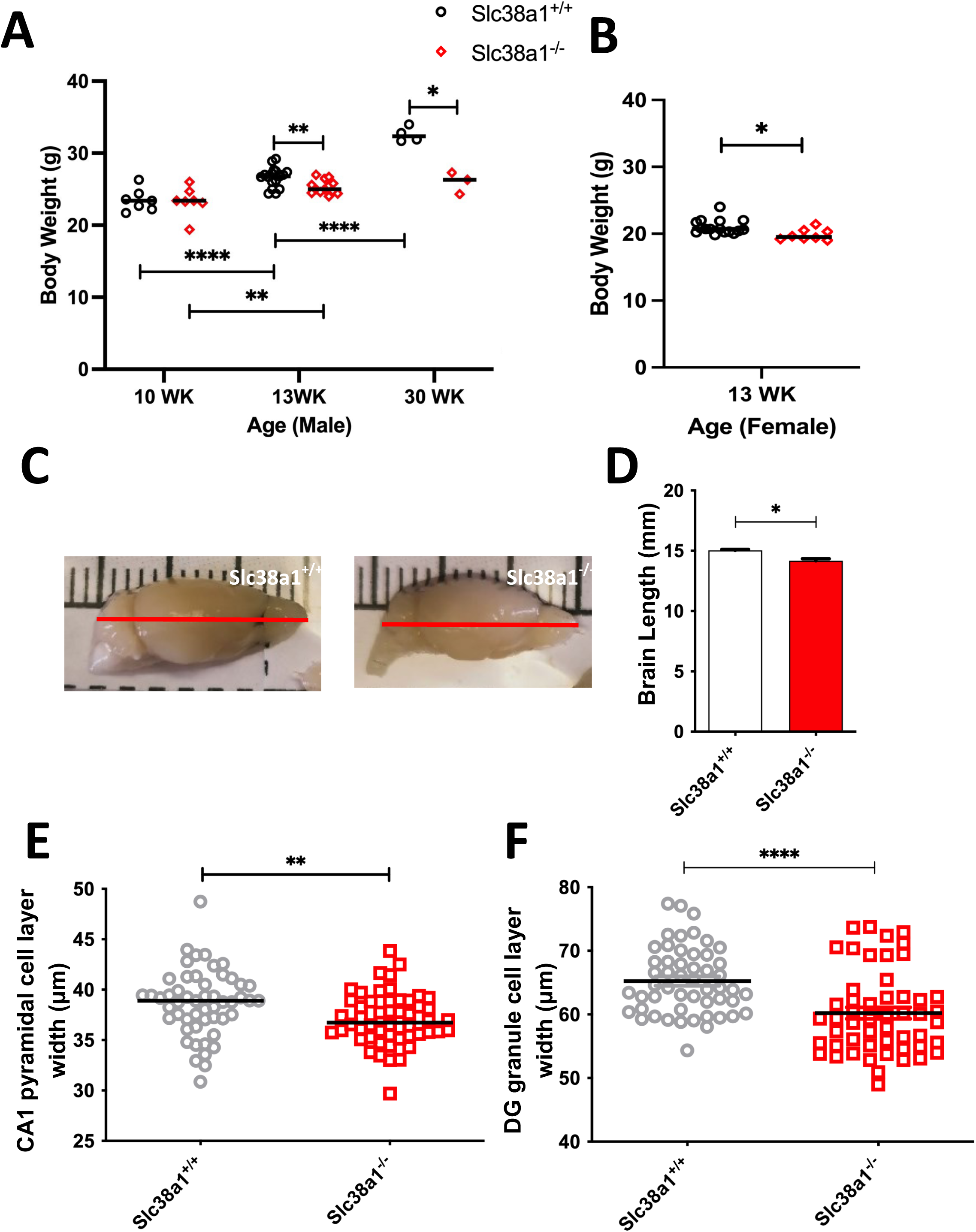
Genetic inactivation of Slc38a1 in mice reduces their brain size and body weight and dilutes numbers of neurons. (A) Body weight of Slc38a1^+/+^ and Slc38a1^−/−^ male mice at three ages (10, 13 and 30 weeks (WK)) were measured. At 10 WKs the two Slc38a1 genotypes have the same weight, but at 13 and 30 WKs of age the Slc38a1^−/−^ mice have significantly lower weight compared to Slc38a1^+/+^ mice. (B) Body weight of Slc38a1^+/+^ and Slc38a1^−/−^ female mice at 13 WKs of age was measured. The female Slc38a1^−/−^ mice have significantly lower weight compared to Slc38a1^+/+^ mice. (C-D) Slc38a1^+/+^ and Slc38a1^−/−^ mice were sacrificed under anesthesia by cervical dislocation followed by harvesting of the brains. The brains were placed on a ruler and the distance from the frontal part to the caudal end of cerebellum was measured and presented in a bar chart. The brain size of Slc38a1^−/−^ mice is significantly smaller compared to brain size of Slc38a1^+/+^ mice. Mean±SEM, N=3. (E-F) Brains were sectioned and the width of the pyramidal cell layer in CA1 and GCL in dentate gyrus was measured. Seven comparable areas were calculated from each of three Slc38a1^+/+^ and Slc38a1^−/−^ mice. Width of both layers are significantly reduced in Slc38a1^−/−^ compared to Slc38a1^+/+^. Male: 10±1 weeks: Slc38a1^+/+^, N=10, Slc38a1^−/−^, N =10; 13±1 weeks: Slc38a1^+/+^, N =20, Slc38a1^−/−^, N =14; 30±1 weeks: Slc38a1^+/+^, N=4, Slc38a1^−/−^, N =3. Female: 13±1 weeks: Slc38a1^+/+^, N=16, Slc38a1^−/−^, N=8. *, P=0.05, **, P=0.01, ***, P=0.001, the unpaired *t*-test statistics was done by Prism GraphPad 9.

We next measured brain length, defined as the distance from the frontal pole of the olfactory bulb to the caudal tip of the cerebellum. This analysis revealed that brains from Slc38a1^−/−^ mice were significantly shorter than those from Slc38a1^+/+^ littermates (Fig. 6C–D), indicating that Slc38a1 disruption affects overall brain development.

To examine potential changes in neuronal architecture, we analyzed the width of two key glutamatergic neuron layers: the CA1 pyramidal cell layer and the dentate gyrus (DG) granule cell layer. Widths were measured from seven anatomically comparable regions per mouse. Both the CA1 and DG layers were found to be significantly thinner in Slc38a1^−/−^ mice compared to Slc38a1^+/+^ (Fig. 6E–F), suggesting a reduction in neuronal number or density in the absence of Slc38a1.

### Slc38a1 inactivation affects neuronal survival and changes cell morphology

To reveal the importance of Slc38a1 in the neuronal development, we isolated primary neuronal cells from both Slc38a1^+/+^ and Slc38a1^−/−^ mouse embryos and cultured them in vitro. Neuronal survival was assessed at day in vitro (DIV) 2 and DIV14. At DIV2, no significant difference in cell number was observed between genotypes (Fig. 7D). However, at DIV14, the cell number of neurons was significantly reduced in the Slc38a1^−/−^ group than Slc38a1^+/+^ group (Fig. 7D). Morphological parameters of cultured neurons were captured and analyzed with the Simple Neurite Tracer (SNT) plugin in ImageJ. The neuronal processes from Slc38a1^−/−^ mice were noticeably shorter than those from Slc38a1^+/+^ mice (Fig. 7A, B). Sholl analysis showed less complexity of neuronal dendrites derived from Slc38a1^−/−^, with lower intersection numbers compared to neurons from Slc38a1^+/+^ mice (Fig. 7C). Although the presence of active Slc38a1 did not influence the branch numbers (Fig. 7E), it did influence average branch length, both first order and second order (Fig. 7F). Altogether, our data imply that active Slc38a1 is essential for neuronal survival and morphology as well as life span of neurons.

**Figure 7.**
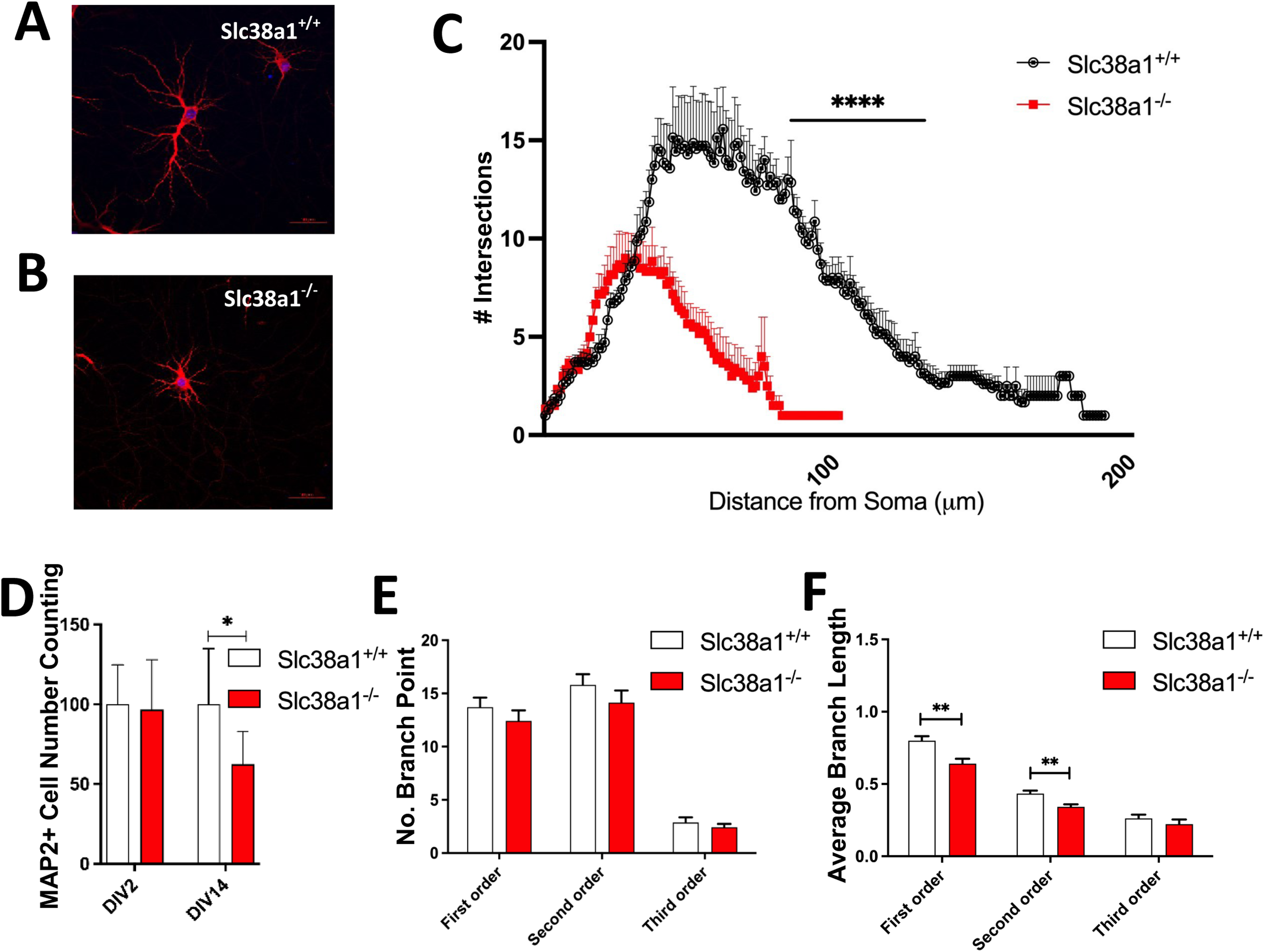
Genetic inactivation of Slc38a1 altered number and morphology of cultured neurons. (A-B) Representative images of Slc38a1^+/+^ and Slc38a1^−/−^ cultured neurons at day *in vitro* (DIV) 14. (C) Sholl analyses of DIV14 cultured neurons were analyzed with Simple Neurite Tracer (SNT) plugin of ImageJ. Slc38a1^−/−^ mice show highly reduced number of intersections and the distance of the intersections from the soma along the processes is decreased (n=21 cells for each point on the curves). (D) Quantification of Slc38a1^+/+^ and Slc38a1^−/−^ cultured neurons show similar cell number at DIV2 but significant reduction at DIV14 in Slc38a1^−/−^ mice. (E-F) The number of branch points are similar in the two genotypes, but the first and second order average branch lengths are significantly reduced in Slc38a1^−/−^ mice (21 cells for both genotypes were analyzed). *, P=0.05, **, P=0.01, ****, P=0.0001, the unpaired *t*-test statistics was done by Prism GraphPad 9, in C using two-way ANOVA.

## DISCUSSION

Our findings demonstrate that Slc38a1 plays a crucial role in embryonic neurogenesis, influencing body weight, brain size, neuronal morphology, and cell survival in adulthood. The robustness of these results is supported by multiple ways: Slc38a1 protein localizes specifically to embryonic neurogenic niches in both mice and rats, and this is corroborated by in situ hybridization and scRNA sequencing showing similar transcript localization in mice (23–25). Additionally, Slc38a1 transcript localization in the adult mouse brain aligns with protein localization in adult rats (22).

### Slc38a1 sustains embryonic neurogenesis through multiple mechanisms and a dysfunction in Slc38a1 affects neuronal volume, morphology and survival

The rapid growth of the embryonic brain demands a high supply of nitrogen and carbon skeletons for DNA and protein synthesis. Glutamine has the highest concentration among amino acids in fetal blood and may fulfill these needs (12, 38). Slc38a1, a high affinity glutamine transporter − is well suited for glutamine uptake in NSCs (14). Glutamine stimulates embryonic NSC proliferation and maintains NSC undifferentiation, partly by stabilizing octamer-binding transcription factor 4 (OCT4) and by stimulating PI3K/Akt and mTOR pathways (7, 39). Consistent with this, theanine − an amino acid and glutamine analogue in green tea − upregulates Slc38a1 and mTOR phosphorylation and promotes proliferation in undifferentiated neuronal progenitor cells (40).

Glutaminase 1, detectable from E11.5 in the mouse brain (41), metabolizes glutamine into glutamate. In humans, glutaminolyses promotes basal progenitor generation, self-renewal and neocortical expansion (8, 9). Fetal glutamate has the highest synthesis rate (3.7 umol/min kg) which exceeds the demand for protein synthesis (38), indicating essential roles in morphogenesis through other mechanisms. Notably, glutamate and GABA act through transmitter-gated receptors before synapse formation, functioning in diffuse, non-synaptic modes (42, 43). These transmitters support self-renewal, proliferation, differentiation, and migration of embryonic stem cells (7, 44, 45). NSCs also establish a glutamate autocrine niche via releasing and responding to the transmitter for their own regulation (46), while the excitatory GABA regulates neurogenesis all the way from NSC proliferation to synaptogenesis (43, 47). Thus, Slc38a1-mediated glutamine transport, and its metabolic products—glutamate and GABA—are essential for embryonic neurogenesis.

Glutamine and its derivatives also serve as morphogens (45), promoting dendritic outgrowth and maturation (30, 42, 48, 49). We now demonstrate broad Slc38a1 expression across both the dorsal pallium and sub-pallium — regions that generate glutamatergic and GABAergic neurons, respectively. Compatible with this, Slc38a1 disruption significantly reduced brain size and neuronal number, and perturbed dendrite development and cellular lifespan, including glutamatergic neurons.

### Slc38a1 in the adult SGZ: indirect modulation via interneurons

Slc38a1 does not co-localize with GFAP^+^, nestin^+^ or Sox2^+^ cells in the SGZ, indicating no direct role in adult neurogenesis. However, GABA released by local PV^+^ and SST^+^ interneurons are critical for successful neural development in adult neurogenesis in the SGZ (30): Intermediate progenitor cells, derived from sub-granular astrocytes/stem cells, can generate immature neurons decorated with GABA-A receptors and be subject to regulation by local GABA release. PV^+^ interneurons regulate activation and proliferation of quiescent adult NSC and make immature synaptic inputs onto proliferating newborn granule cells (50, 51). SST^+^ interneurons also establish synaptic contacts with granule cells, but they are restricted to granule cell dendrites (35). Through these mechanisms, GABA signaling shapes dendritic development and synaptogenesis (30). The structural and functional diversity of PV+ and SST+ interneurons enables precise temporal and spatial control of GABAergic modulation.

### Slc38a1 is enriched in ependyma, but not linked to adult neurogenesis

Slc38a1 is highly expressed in adult ventricular ependymal cells, consistent with single cell transcriptomics data (52). However, Slc38a1 does not co-localize with nestin^+^ or GFAP^+^ ependymal cells, contradicting a role in adult neurogenesis. Frisen et al. reported that ependymal cells have NSC properties able to rapidly generate neurons, which migrate to the olfactory bulb or convert to astrocytes for scar formation upon spinal cord injury (53, 54). This has been disputed by several other research teams: Alvarez-Buylla et al. show that adult ependymal cells are post-mitotic derived from radial glial cells and that NSC B1 are displaced from VZ into SVZ by ependymal cells (55–57). Biernaskie’s group also point out that ependymal cells and adult NSC B1 cells do have similar embryonic origins, however, they are transcriptionally distinct, and they fail to exhibit stem cell functions (58). We have earlier shown that the homologues Slc38a3 (aka SN1 and SNAT3) is highly expressed in ciliated ependyma cells (34). We have also shown that Slc38a3 and Slc38a1 work in concert to transport glutamine intercellularly or across epithelial cells (14, 16, 59). Slc38a1, together with Slc38a3, in the ependyma cells is therefore more likely to shuttle glutamine from ventricles to or across ependymal cells to the SVZ for metabolism, and may possibly contribute indirectly to adult neurogenesis, e.g., through GABAergic interneurons (30).

## Conclusion

The unique kinetical properties, regulation and affinity for glutamine allow Slc38a1 to efficiently accumulate glutamine into embryonic neurogenic niches, thereby supplying rapidly growing cells with ample glutamine for the formation of building blocks and energy. The significance of Slc38a1 in the neurogenesis of both glutamatergic, GABAergic and glial cells is bolstered by our finding of enriched Slc38a1 protein as well as RNA, both by *in situ* hybridization and scRNAseq analyses, in the dorsal pallium and sub-pallium. Slc38a1 is highly regulated in time and space: e.g., at E15.5, Slc38a1 in the forebrain is highest in the VZ, while enriched in SVZ in the hindbrain, which is more mature. In the adult, the neurogenic cells in the SGZ are devoid of Slc38a1, while Slc38a1 is enhanced in dentate gyrus local PV^+^- and SST^+^-interneurons, which connect to and regulate the neurogenic cells in the SGZ. Finally, Slc38a1 maintains neuronal survival and number, brain size and cell morphology. Dysfunction of Slc38a1 is likely to contribute to dendrite pathology and neuropsychiatry (unpublished data). Encouragingly, transplantation of medial ganglionic eminence (MGE) progenitors into the hippocampus yields interneurons that restore behavioral deficits such as impaired learning and hyperactivity (65). These findings suggest that progenitor cell therapies targeting Slc38a1 dysfunction may hold promise for treating neuropsychiatric diseases in the future.

## Acknowledgements

Thanks to Professor Torkel Hafting for WFA antibody and to Dr. Chencheng Wang for Sox2 antibody.

