## Supplementary information for "The glutamine transporter Slc38a1 is widely expressed in the embryonic neurogenic niches and impacts neuronal volume, survival, and morphology"

**Abbreviated title: Slc38a1 in Neurogenesis**

**Authors:** Daxin Wang[1], Marivi Nabong Moen[1], Diana Domanska[2,3], Muhammad Zahoor[1], Wenjing Cai[1], Ragnhild Elisabeth Heimtun Paulsen[4], Tor Paaske Utheim[5], Frode Lars Jahnsen[2,6], Jon Storm-Mathisen[1] and Farrukh Abbas Chaudhry[1,2]

[1]Department of Molecular Medicine, Institute of Basic Medical Sciences, University of Oslo, Norway

[2]Department of Pathology, Oslo University Hospital – Rikshospitalet, Norway

[3]Faculty of Mathematics and Computer Science, University of Warmia and Mazury in Olsztyn, Olsztyn, Poland

[4]Section for Pharmacology and Pharmaceutical Biosciences, University of Oslo, Norway

[5]Department of Plastic and Reconstructive Surgery, Oslo University Hospital – Rikshospitalet, Norway

[6]Institute of Clinical Medicine, University of Oslo, Norway

**Corresponding author:** Farrukh Abbas Chaudhry, Department of Molecular Medicine, Institute of Basic Medical Sciences, University of Oslo, Norway.

### MATERIALS AND METHODS

#### Animals

The animal handling and all experiments were conducted in strict accordance with the recommendations described in the Norwegian Animal Welfare act and the European Union Directive on the protection of Animals used for Scientific Purposes (2010/63/EU). All experiments have been approved by the Norwegian Food Safety Authority (FOTS application numbers 9200, 21009 and 30120) and reported in accordance with reporting on In Vivo Experiments (ARRIVE; du sert 2020). Mice were housed in a room with fixed 12-hour light/dark cycle, temperature set to 22 ± 1 °C with relative humidity at 50 ± 10%. The animals were stalled in groups of up to 5 mice in cages made of Makrolon GreenLine cages (Sealsafe Plus GM500 or GM900, Buguggiate, Italy). The cages were ventilated with 100% fresh air at all times except during experiments. The cages were enriched with paper for nest-building, a wooden stick, a paper roll or a small plastic house and a running wheel. 3 Rs (Replacement, Reduction and Refinement) principles were applied in the experiments design. Autoclaved food (RM3 from Special Diets Service (UK) and water were provided ad libitum.

Animals included in the study were of both sexes and were housed in the animal facility, Section of Comparative Medicine (KPM), Institute of Basic Medical Sciences, University of Oslo. The *Slc38a1*$^{-/-}$ mouse model used in these studies has previously been rigorously characterized phenotypically (1),(2). Animals used for the study were backcrossed in to C57BL/6J for 10 generations. Mice that were wild type (*Slc38a1*$^{+/+}$) or genetically inactivated for *Slc38a1* (*Slc38a1*$^{-/-}$) were all originating from the same breeding colony. The mice were earmarked for identification at 4 weeks of age. The biopsies taken during earmarking were used for genotyping. Female animal vaginal plug check service was provided by the experienced staff at KPM. The day when females appear vaginal plugs was defined as Embryonic day 0.5 (E0.5).

Age and weight of all Slc38a1$^{+/+}$ and Slc38a1$^{-/-}$ mice used in quantitative western blotting, immunofluorescence labeling experiments and *in situ* hybridization were noticed and/or measured before termination.

**Quantitative western blotting**

Quantitative western blotting was performed as detailed in our previous papers (3);(4) with some modifications. Briefly, three pairs of age- and sex-matched Slc38a1$^{+/+}$ and Slc38a1$^{-/-}$ mice at specific developmental stages (i.e., pregnant mice for E15.5 embryo and P1, P7, P14, P20 and adult mice or pregnant rats for E15.5 and adult rat) were anaesthetized and decapitated, brains were rapidly dissected out, frozen in dry ice and stored at -80°C. The tissue was homogenized in RIPA (10mM Tris-HCl, pH 8.0; 1mM EDTA; 0.5mM EGTA; 1% Triton X-100; 0.1% Sodium Deoxycholate; 140mM NaCl; proteinase inhibitor; phosphatase inhibitor, dilute with dH2O) and mixed with 0.1% SDS before use. 10% Criterion™ TGX™ Precast Midi Protein Gel was used for protein separation and Trans-BlotTurbo Transfer System RTA Transfer Kits was used for transferring to the blots according to the manufacturer's instruction. 5% skim milk in TBST (0.05 M Tris-HCl, pH 7.4, 0.9% sodium chloride, and 0.1% Triton) was applied to block unspecific staining. The blots were incubated with primary antibodies of interest in 3% skim milk in TBST overnight at 4°C. After rigorously washing, the blots were incubated with secondary antibody conjugated with horseradish peroxidase in 3% skim milk in TBST at room temperature (RT) for 1 hour. After rinsing with TBST 3x 10 min, SuperSignal™ West Pico PLUS Chemiluminescent Substrate and ChemiDoc Imaging Systems were used to capture the images. The normalization to GAPDH bands of the same blots was applied to minimize loading errors. The band signal intensity was measured by BioRad Image Lab 6.0 software.

**Immunofluorescence staining**

Age- and sex-matched Slc38a1$^{+/+}$ and Slc38a1$^{-/-}$ mice and rats at specific developmental stages (i.e., pregnant mice for E15.5 and E18.5 embryo and P1, P7, P14, P20 and adult mice or pregnant rats for E15.5 and E18.5 embryo) were anaesthetized and transcardially perfusion-fixed with 4% fresh made paraformaldehyde (PFA). Brains were rapidly dissected out, the size of the left or right hemisphere of the brains measured on a paper with scale of ruler followed by immersion in 4% PFA overnight. However, for embryonic brain, the embryo head skin was removed, a midline ventral incision was made to allow PFA immerse the brain tissue. The hemispheres were then treated with 10% sucrose solution in the first day, 20% sucrose solution in the second day and 30% sucrose solution in the third day. After complete sinking of the brain hemispheres into the bottom of the tube with 30% sucrose solution, they were frozen with dry ice followed by embedding with Neg-50 Frozen Section Medium (Nerliens Meszansky AS, Cat. No. SHA6502). Coronal or sagittal sections with 18μm thickness were cut by a cryostat machine. The sections were rinsed with phosphate-buffer saline (PBS) 3 x 10 min followed by incubation in blocking buffer containing 5% goat or donkey serum, 1% BSA and 0.5% Triton in TBS for 1 hour at room temperature (RT). Primary antibodies of interest were diluted in TBS with 3% goat or donkey serum, 1% BSA and 0.5% Triton, and sections incubated in this solution overnight at 4°C. Following 3 washes with PBS, the sections were incubated in diluted Alexa Fluor™ secondary antibodies for 2 hours at RT. After washing with PBS, 5μl 4',6-diamidino-2-phenylindole (DAPI) was added on the section for 10 minutes. ProLong™ gold Antifade Mountant was applied to get a better fluorescence result.

***In situ* hybridization with RNAscope probe in embryonic and adult brain**

Sample preparation: Age- and sex-matched Slc38a1$^{+/+}$ and Slc38a1$^{-/-}$ mice were anesthetized followed by a transcardial perfusion with 4% PFA. After perfusion, brains were rapidly dissected out and immersed into 4% PFA overnight. The brains were then incubated in increasing concentrations (10-30%) of sucrose for at least 3 days until they sunk into the bottom of the container. The brains were then frozen with Neg-50 Frozen Section Medium and sectioned by cryostat with thickness of 14 µm.

Chromogenic In situ hybridization (ISH) was performed for Slc38a1 (RNAscope® Probe - Mm-Slc38a1-C1, Mus musculus solute, Cat. No: 1143671-C1, Bio-techne) using The RNAscope® 2.5 High Definition(HD)- BROWN Assay kit, Cat. No. 322300, Bio-techne) in accordance with the manufacturer's instructions. The sections were mounted with Eukitt and dried at RT overnight.

Data analysis was performed using images captured with 20× and 40× objective lenses. Each brown dot represents a single RNA transcript. Images were taken from entire brain sections or specific brain regions (sub-ventricular zone, dentate gyrus). For analysis, at least three randomly selected 40× fields per region were converted to 8-bit in Fiji/ImageJ. The color threshold was adjusted and used the same settings across groups. Particle counts (RNA transcript numbers) were quantified, and comparisons between groups were made using a non-parametric t-test in GraphPad Prism 9.

**Hippocampal cornu ammonis 1 (CA1) and granule cell layer (GCL) width measurements**

A series of sagittal 18 µm thick sections were made by a cryostat through the whole hemisphere. At least 3 comparable sections were chosen from these sections to stain with DAPI. To measure the width of GCL of hippocampus, the border of GCL was defined as the border of the last nuclear contact to the main layer. To avoid variation, only the upper GCL in dentate gyrus (DG) was measured. Seven randomized and comparable regions of interest were chosen from DG and CA1 for the measure of the width. Adobe PhotoShop CS6 was applied to assist the measurement. A non-parametric t-test was performed by GraphPad Prism 9.

**Neuron survival counting**

Primary neuron cell culture is a standardized protocol in our lab. Briefly, embryonic (E18.5) brains from Slc38a1$^{+/+}$ and Slc38a1$^{-/-}$ mice were carefully dissected and placed into a dish with complete medium. Brain meninges were carefully removed, brain cortex cut into small pieces and trypsin solution was added to help digesting the tissues into single cells. To reduce the sticky cell suspension, DNAse was used for digesting the DNA released from the brain tissue. At least 60,000 cells were plated into each well on cover slips of 24-well plate. The primary cells were incubated at 37°C and 5% $CO_2$ and the cell medium changed every 3rd days. The primary cortex neuronal cultures matured after 2 weeks' incubation. The primary neurons were then incubated with antibody against MAP2 in 4°C overnight and completed as described for Immunofluorescence staining above. MAP2 is considered mainly expressed by the dendrites of neuronal cells. Hence, a cell with both MAP2 and DAPI was counted as a mature neuron here. Seven randomized confocal microscope views (20x) were captured and the cells counted manually and blindedly by the counting tool of Adobe PhotoShop CS6. Only cells with the whole nuclear inside were counted. A non-parametric t-test was performed by GraphPad Prism 9.

**The Sholl analysis of dendrites**

The Sholl analysis were performed by the assistance of Fiji plugin Simple Neurite Tracer and measured according to the manual from imagej.net. The starting radius was set at the beginning of the longest dendrites of each cell and the ending radius and radius step size were set as recommended or default. 21 cells of 14-days *in vitro* (DIV14) with MAP2$^{+}$ at a magnification of 40x from each genotype were randomly selected in this study and the intersections, branch numbers and lengths were calculated. Non-parametric t-test and Two-way ANOVA was performed by GraphPad Prism 9.

**Membrane trafficking of Slc38a1 in Hela Cells**

Human origin Hela cells were seeded on glass coverslips and then transfected with Slc38a1-GFP plasmid for 24 hours, using TransIT®-LT1 Transfection Reagent (Mir2304, Mirusbio). The cells were then fixed with 4% PFA for 6 minutes at room temperature and then incubated with 50mM of NH4Cl for 8 minutes for quenching. Next, the cells on cover glass were washed with PBS and then permeablized with 0,25% triton X-100 for 8 minutes and followed by through washing with PBST (0,05 tween 20 in PBS), three times.  Cells were blocked with 5% BSA in PBST for 1 hour at room temperature, before incubating with primary antibodies (dilution factor 1:1000) at room temperature for 1 hour in incubation buffer (1% BSA in PBST). The cells were washed wtih PBST three times, followed by incubating with secondary antibodies (dilution factor 1:1000) at room temperature for 1 hour. The cover glasses were washed with PBST (3X) and then breifly rinsed with Mili-Q water before embedding in polyvinyl alcohol mounting medium (Sigma, catalog # 10981). The images were acquired on LSM700 confocal microscope, using PlanAphochromat 63x/1.40 oil Ph3 M 27 objective. The detail of antibodies is as followed along with catalog numbers.

**Figure S1. Slc38a1 protein expression level in the embryonic rat brain is comparable to the expression level in the adult rat brain**

(A) Whole rat brain extracts from the embryonic day 15.5 (E15.5) and adult were electrophoretically separated in SDS-PAGE followed by immuno-labeling for Slc38a1. Anti-Slc38a1 antibody reveals a single specific band at ~50 kDa, both in the embryonic and in the adult brains.

(B-E) E15.5 embryonic rat brain sections were tripled labeled for Slc38a1 (green), cell-specific markers (red) and DAPI (blue).

(B) Merged images of Slc38a1, Sox2 and DAPI staining show that Slc38a1 labeling surrounds $Sox2^+/DAPI^+$ (*) as well as $Sox2^-/DAPI^+$ ($^{\#}$) stained cell nuclei. The marked area is enlarged and shown in Figure 1F.

(C-D) In the sub-pallium, Slc38a1 staining surrounds both $Sox2^+/DAPI^+$ (*) and $Sox2^-/DAPI^+$ ($^{\#}$) nuclei in the ventricular zone (VZ) and/or sub-ventricular zone (SVZ), both in the lateral ganglionic eminence (LGE) and the medial ganglionic eminence (MGE).

(E) In the hindbrain, Slc38a1 staining is primarily limited to the SVZ where $Scl38a1^+$/DAPI cells (*) appear in between Nestin labeled cell processes (arrow head).

Nestin is a neuroepithelial stem cell marker, while Sox2 is a marker for neuroepithelial stem cell and progenitor cells. DAPI visualizes nuclear DNA. Scale bars represent 20 um (B-E). LV: Lateral ventricle.

**Figure S2. In the adult brain, Slc38a1 protein does not co-localize with $nestin^+$- or $GFAP^+$ cells in sub-granular zone and ependymal layer, but localizes in $parvalbumin^+$ interneurons**

(A-D) Brain tissue from adult mice were labeled for Slc38a1 (green), cellular markers (red) and DAPI (blue) and investigated for a role of Slc38a1 in adult neurogenesis. (A) Slc38a1 staining does not co-localize with Nestin$^{+}$ cells in the hippocampal dentate gyrus in adult Slc38a1$^{+/+}$ mice, expect for some bleed-through.

(B-C) Slc38a1 is expressed in most ependymal cells lining the lateral ventricle (LV) of Slc38a1$^{+/+}$ mice (arrow), but co-localize not with GFAP$^{+}$ ependymal cells or their processes in the sub-ventricular zone (arrowhead). No staining for Slc38a1 was detected in the ependymal cells in Slc38a1$^{-/-}$ mice.

(D) Double labeling experiments with antibodies against Slc38a1 and *Wisteria floribunda* agglutinin (WFA), a lectin binding to *N*-acetylgalactosamine, which is a component of perineuronal nets in the extracellular matrix of PV$^{+}$ interneurons, reveal co-localization in scattered cells in the hilus of the DG. Scale bars represent 20 um (A-C), 10 μm (D).

**Figure S3. Membrane trafficking of Slc38a1 suggests tight on-demand regulation of Slc38a1 at the plasma membrane**

The intracellular landscape of Slc38a1. Hela cells transfected with Slc38a1 for 24 hours were stained/imaged for endomembrane compartments markers; Plasma membrane A), RTN3 B), TGN46 C), GM130 D), EEA1 E) and CD63 F). Intensity profiles (insets in C-F, taken from cells other than those shown in the picture) of the two channels are overlayed to demonstrate co-localization in a different way than colour mixing to yellow.

(A) Labeling for Slc38a1 is detected on the plasma membrane (PM; arrow).

(B) Slc38a1 staining is accumulated in the peri-nuclear region and co-localizes partially with RTN3, a marker for endoplasmic reticulum.

(C-D) Slc38a1 co-localizes (yellow) with a marker for the trans-Golgi network (TGN46) and by intensity profile, while Slc38a1 and cis-Golgi (GM130) staining and the channel overlay do not lend support for extensive co-localization.

(E-F) Slc38a1 colocalizes well with markers for early endosomes (EEA1) and with a marker for late endosomes (CD63).

**Figure S4. High level of Slc38a1 transcript in several embryonic mouse brain areas and in select cells in the adult**

***A***, Coronal sections were made of E18.5 and adult mouse forebrain, labeled with complementary RNA (anti-sense) for Slc38a1 and the localization investigated. There are extensive probe particles for Slc38a1 transcript throughout the section. The areas limited by the rectangles are magnified in Figure 4A, 4C and 4D. ***B-C***, High number of Slc38a1 probe particles are enriched in the embryonic medial mammillary nucleus (Mmn) and lateral hypothalamus (LHy). ***D***, Strong labeling with Slc38a1 antisense RNA probes is seen in some individual cells in the hilus (Hi) of the hippocampal dentate gyrus (DG; arrows). The granule cells are also moderately stained by Slc38a1 RNA probe (arrowheads). ***E,*** In the adult lateral ventricular (LV) wall, strong labeling by Slc38a1 probe is seen in the ependymal layer (Ep), while most of sub-ventricular zone (SVZ) do not show labeling for Slc38a1 except for some scattered cells (red arrow). ***F***, In the wall of 4th ventricle, strong labeling for Slc38a1 transcript is enriched in Ep cells (red arrow) and some scattered cells in the SVZ (white arrow). The rest of the sub-ventricular zone (white arrowhead) and the choroid plexus (CP; red arrowhead) are devoid of any labeling.
