## Supplementary figures for "The glutamine transporter Slc38a1 is widely expressed in the embryonic neurogenic niches and impacts neuronal volume, survival, and morphology"

### Figure S1

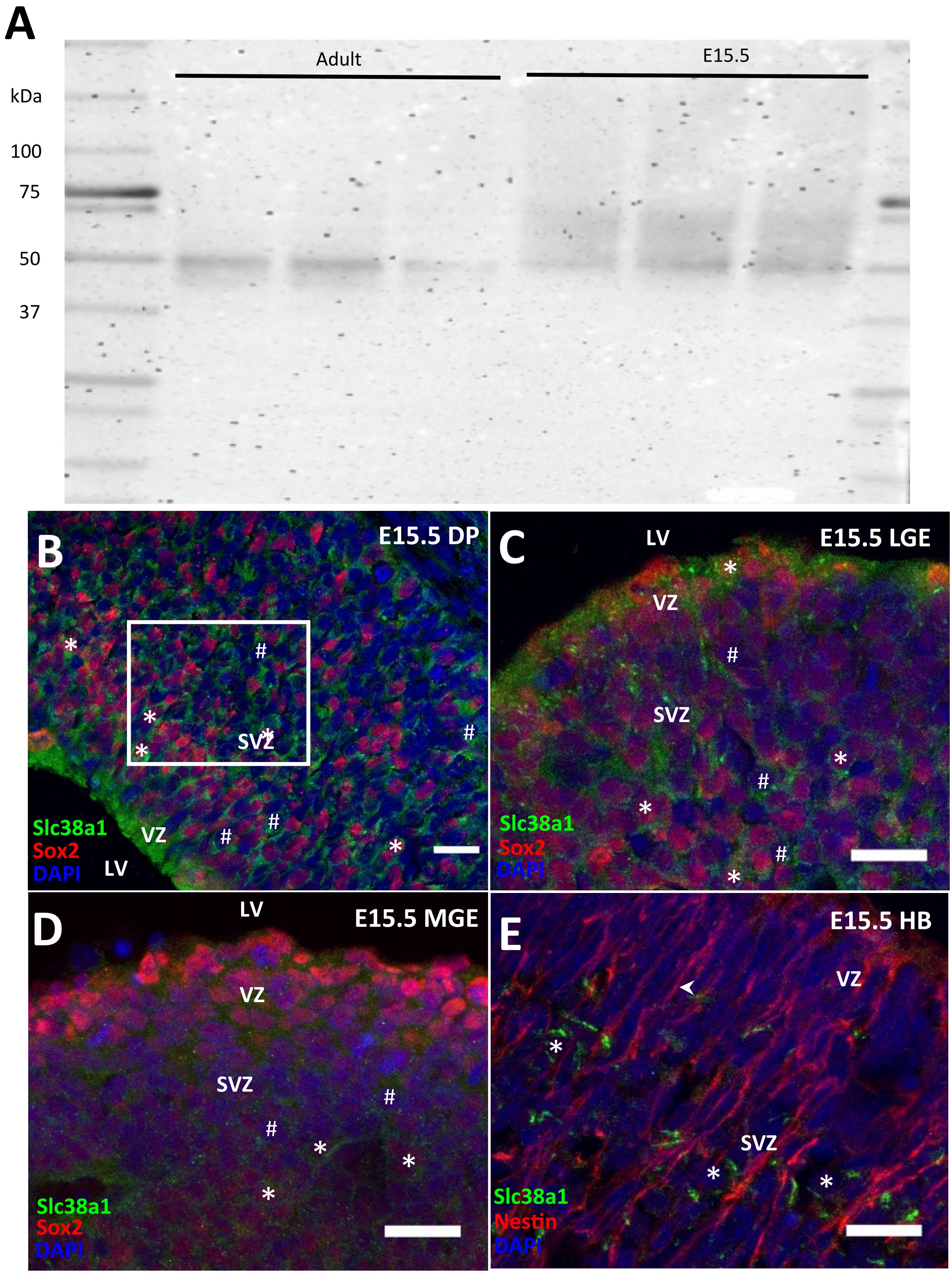

A
kDa
100
75
50
37
Adult
E15.5
B
E15.5 DP
SVZ
VZ
LV
Slc38a1
Sox2
DAPI
C
LV
E15.5 LGE
VZ
SVZ
Slc38a1
Sox2
DAPI
D
LV
E15.5 MGE
VZ
SVZ
Slc38a1
Sox2
DAPI
E
E15.5 HB
VZ
SVZ
Slc38a1
Nestin
DAPI

### Figure S2

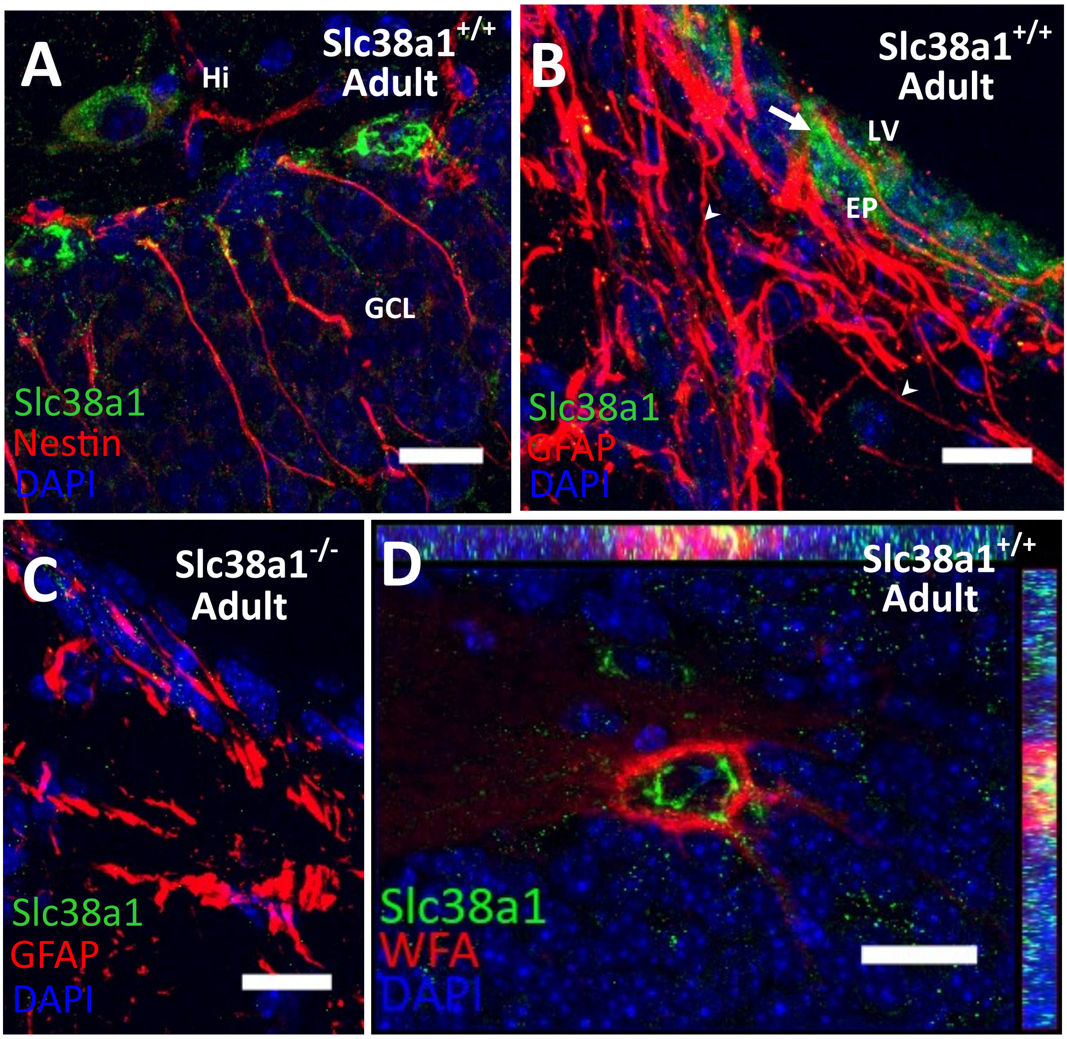
A
Slc38a1+/+
Adult
Hi
GCL
Slc38a1
Nestin
DAPI
B
Slc38a1+/+
Adult
LV
EP
Slc38a1
GFAP
DAPI
C
Slc38a1-/-
Adult
Slc38a1
GFAP
DAPI
D
Slc38a1+/+
Adult
Slc38a1
WFA
DAPI

### Figure S3

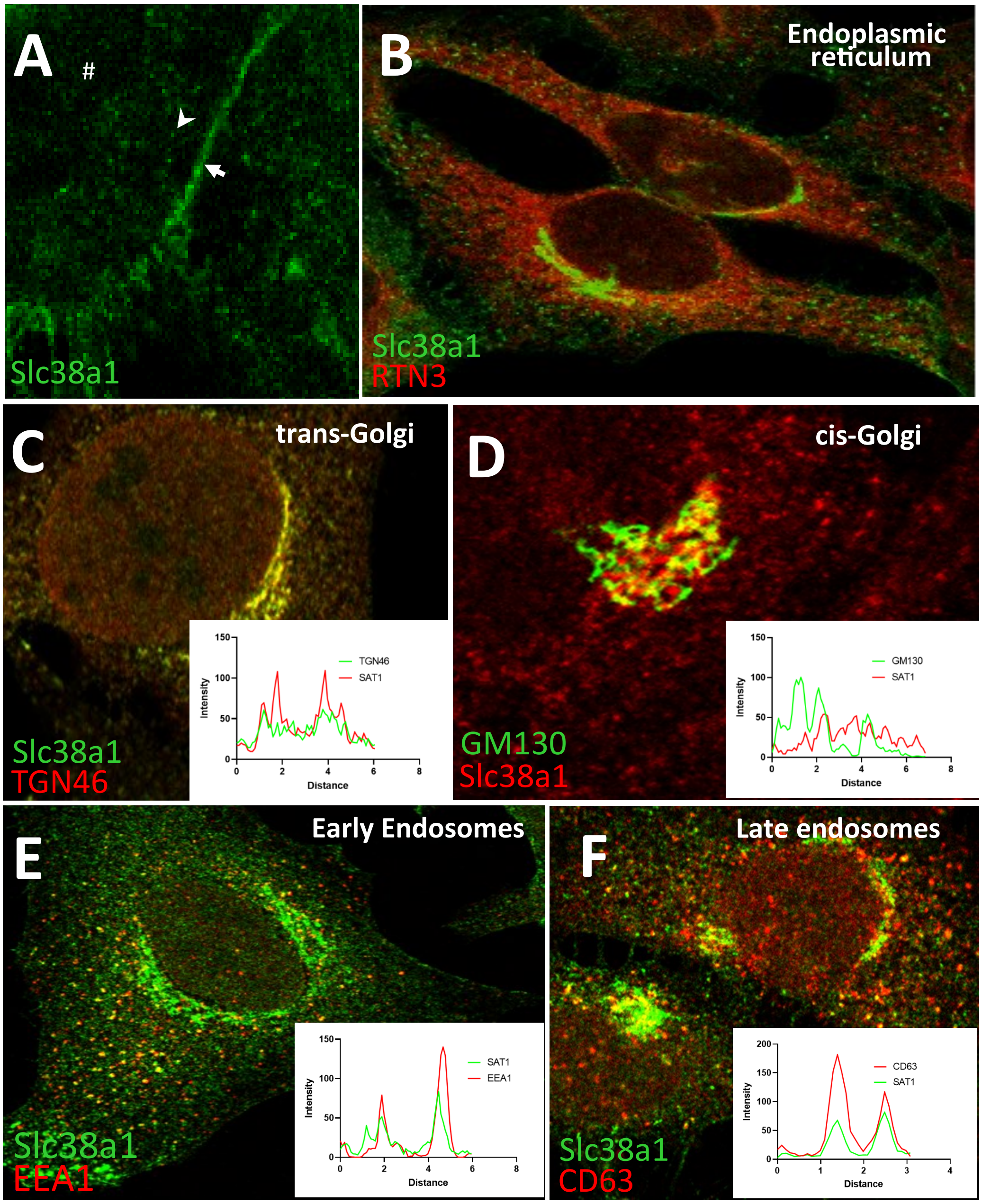

A
#
Slc38a1
B
Endoplasmic reticulum
Slc38a1
RTN3
C
trans-Golgi
Slc38a1
TGN46
TGN46
SAT1
Intensity
Distance
D
cis-Golgi
GM130
Slc38a1
GM130
SAT1
Intensity
Distance
E
Early Endosomes
Slc38a1
EEA1
SAT1
EEA1
Intensity
Distance
F
Late endosomes
Slc38a1
CD63
CD63
SAT1
Intensity
Distance

### Figure S4

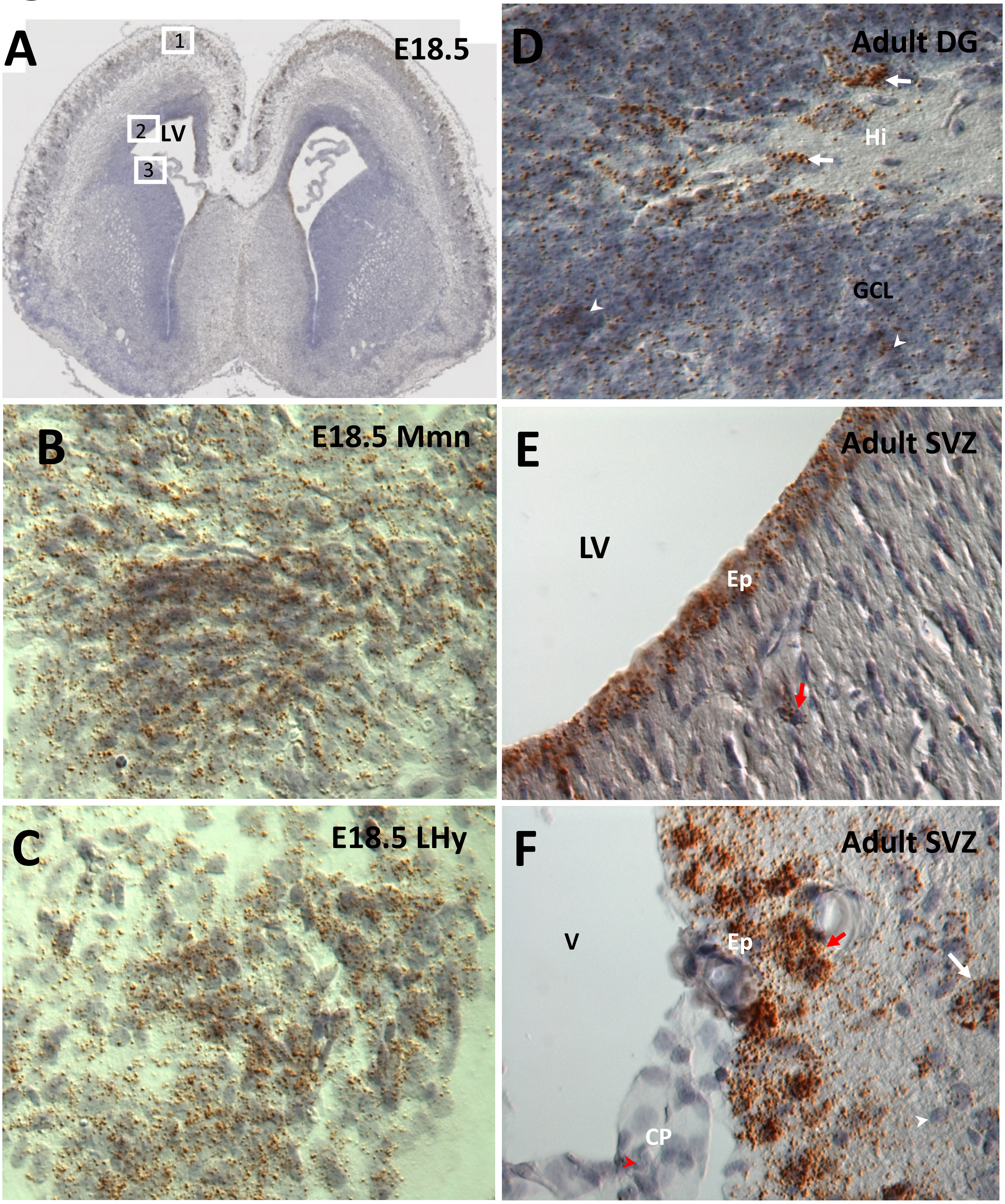
A
E18.5
1
2
LV
3
B
E18.5 Mmn
C
E18.5 LHy
D
Adult DG
Hi
GCL
E
Adult SVZ
LV
Ep
F
Adult SVZ
V
Ep
CP

### Supplementary Table 1

| Name | supplier | Cat. No |
|---|---|---|
| Trans-Blot Turbo 5x Transfer Buffer | BioRad | 10026938 |
| 10% Criterion™ TGX™ Precast Midi Protein Gel | BioRad | 5671034 |
| SuperSignal™ West Pico PLUS Chemilumi-nescent Substrate | ThermoFischer | 34580 |
| Neurobasal™ Medium | Fisher Scientific | 21103049 |
| B27 Supplement | Fisher Scientific | 11530536 |
| Prolong Diamond Antifade Mountant | Fisher Scientific | P36970 |
| ETHANOL 96% F 230 | Univar Solutions AS | 55023 |
| Glacial Acetic Acid | VWR Intl. AS | 20104 |
| Trizma® hydrochloride | Merck-Sigma | T3253 |
| Trizma® base | Merck-Sigma | T1503 |
| Sodium chloride solution | Merck-Sigma | S5150 |
| Tween-20 | Merck-Sigma | P2287 |
| Skim Milk powder | Merck-Sigma | 70166 |
| Ponceau S | Merck-Sigma | P3504 |
| Sucrose | Merck-Sigma | S0389-500G |
| PhosSTOP | Merck-Sigma | 4906837001 |
| cOmplete ULTRA Tablets, Mini EDTA free, EASY pack | Merck-Sigma | 5892791001 |
| Sodium deoxycholate | Merck-Sigma | D6750 |
| Paraformaldehyde | Merck-Sigma | P6148 |
| Triton X-100 | Merck-Sigma | X100 |
| Bovine serum albumin | Merck-Sigma | A9647 |
| DAPI | Merck-Sigma | D9542 |
| EDTA | Merck-Sigma | EDS |
| RNAscope™ 2.5 HD Reagent Kit-BROWN | BioTechne | 322300 |
| Neg-50 Frozen Section Medium | Nerliens Meszansky AS | SHA6502 |
| Eukitt | Fluka | 3989 |

### Supplementary Table 2

| Name | supplier | Cat. No |
|---|---|---|
| Anti-Slc38a1 | Home-made | none |
| Anti-Sox2 | Santa cruz | sc-365823 |
| Anti-RTN3 | Santa cruz | sc-374599 |
| Anti-NeuN | Millipore | MAB377 |
| Anti-Map2 | Millipore | MAB5543 |
| Anti-Somatostatin | Millipore | MAB354 |
| Anti-Parvalbumin | Millipore | MAB1572 |
| Anti-Gapdh | Millipore | MAB374 |
| Anti-Gfap | Sigma | G3893 |
| Anti-Wisteria floribunda agglutinin , Biotin | Sigma | L1516 |
| CD63 | Sigma | HPA010088 |
| Anti-Gad67 | Abcam | ab26116 |
| Anti-TGN46 | Abcam | Ab50595 |
| Anti-Nestin | Novusbio | NB100-1604 |
| Anti-EEA1 | BD Biosciences | 610457 |
| Anti-GM130 | Cell Signaling | 12480S |
| Alexa Fluor 488 goat anti-rabbit IgG | Thermo Fischer | A11034 |
| Alexa Fluor 555 donkey anti-mouse IgG | Thermo Fischer | A31570 |
| Alexa Fluor 555 goat anti-chicken IgG | Thermo Fischer | A21437 |
| Alexa Fluor 555 donkey anti-rabbit IgG | Thermo Fischer | A31572 |
| Alexa Fluor 488 Goat anti-Rat IgG | Thermo Fischer | A11006 |
| Streptavidin, Alexa Fluor™ 594 conjugate | Thermo Fischer | S11227 |
| Anti-rabbit IgG Horseradish Peroxidase | Thermo Fischer | 31460 |
| Anti-mouse IgG Horseradish Peroxidase | Thermo Fischer | 31430 |
